# Homeostatic control of an iron repressor in a GI tract resident

**DOI:** 10.1101/2023.01.30.526184

**Authors:** Yuanyuan Wang, Yinhe Mao, Xiaoqing Chen, Kaiyan Yang, Xinhua Huang, Lixing Tian, Tong Jiang, Yun Zou, Xiaoyuan Ma, Chaoyue Xu, Zili Zhou, Xianwei Wu, Lei Pan, Huaping Liang, Changbin Chen

## Abstract

The transition metal iron plays a crucial role in living cells. However, high level of iron is potentially toxic through the production of reactive oxygen species (ROS), serving as a deterrent to the commensal fungus *Candida albicans* for colonization in the iron-rich gastrointestinal (GI) tract. We observe that the mutant lacking an iron-responsive transcription factor Hap43 is hyper-fit for colonization in murine gut. We demonstrate that high iron specifically triggers multiple post-translational modifications (PTMs) and proteasomal degradation of Hap43, a vital process guaranteeing the precision of intestinal ROS detoxification. Reduced levels of Hap43 lead to de-repression of antioxidant genes and therefore alleviate the deleterious ROS derived from iron metabolism. Our data reveal that Hap43 functions as a negative regulator for oxidative stress-adaptation of *C. albicans* to gut colonization and thereby provide a new insight into understanding the interplay between iron homeostasis and fungal commensalism.

**Importance:** Iron homeostasis is critical for creatures. *Candida albicans* is one of the major commensals in the GI tract where is iron-replete environment. Transcriptional factor Hap43 was believed to repress iron utilizations genes in iron-depleted conditions for decades. However, the mystery in iron-replete conditions of Hap43 has never been uncovered. We discovered that reduced levels of Hap43 via phosphorylation-dependent nuclear export, followed by proteosome-mediated protein degradation, leads to de-repression of downstream antioxidant genes and promote its colonization in GI tract. We propose that *C. albicans* has a strict detoxification process to ensure its survival, which has important implications for understanding how the fungi survives in the mammalian host.

## Introduction

Iron is an essential element required for the viability of virtually all organisms (Andrews, 2008). This transition metal acts as an enzyme cofactor, predominantly in electron transfer and catalysis, and therefore contributes to numerous metabolic processes, in particular energy generation, oxygen transport and DNA synthesis. However, excess of iron is toxic and potentially fatal, primarily because reactive oxygen species (ROS), including hydroxyl radicals (OH^·^), superoxide (O2^·^) and H_2_O_2_, are generated through the iron-catalyzed Haber-Weiss/Fenton reaction and causes cell damage and death (Pierre et al., 2002). The mammalian gut is considered as an iron-rich environment where large amounts of dietary iron (e.g. ∼0.27 mM per day in humans) are regularly present in the colonic lumen and remain unabsorbed (Miret et al., 2003). Interestingly, previous studies have found that the concentration of iron in feces of British adults regularly consumed a standard Western diet and of weaning infants fed with complementary foods could reached to an average value of 100 μg/g wet weight feces (∼1.8 mmol), and this level is actually much higher than the minimal concentration (∼0.4 mmol) required for intestinal bacterial species (Lund et al., 1998, 1999; Pizarro et al., 1987). Hence, it is very likely that the increased level of unabsorbed iron would aggravate the status of oxidative stress in the gastrointestinal (GI) tract, providing a detrimental impact on the growth of resident microbial commensals. Indeed, excessive ROS levels in this iron-replete niche were found to enhance cellular toxicity, reflected by oxidative damage to proteins, lipids and DNA, and therefore restrict the growth and proliferation of colonized microorganisms (Dixon & Stockwell, 2014; Schieber & Chandel, 2014).

The potent redox capability of iron requires that microbes carefully respond to and regulate environmental iron levels and distribution (Barber & Elde, 2015). Several examples of iron detoxification have been described in bacterial species. For example, both *Escherichia coli* and *Salmonella Typhimurium* developed effective iron efflux systems to decrease intracellular accumulation of free iron and prevent iron stress (Frawley et al., 2013; Grass et al., 2005). In similar, eukaryotic microbes like fungi also decreased the labile iron pool to prevent formation of deleterious hydroxyl radicals through the vacuolar and siderophore-mediated iron storage (Gupta & Outten, 2020; Singh et al., 2007). However, studies investigating mechanisms employed by gut microbes to detoxify iron-mediated ROS production are relatively limited.

*Candida albicans* is a major opportunistic fungal pathogen of humans, capable of causing mucosal diseases with substantial morbidity and life-threatening bloodstream infections in immunocompromised individuals (da Silva Dantas et al., 2016; Noble & Johnson, 2007). Importantly, this fungus is also the most prevalent fungal species of the human microbiota and acts as a commensal to effectively colonize many host niches, particularly the GI tract (Kumamoto et al., 2020). Our previous studies have demonstrated that the acquisition and storage of iron in *C. albicans* was effectively regulated by a complex and effective regulatory circuit, which consists of three iron-responsive transcription regulators (Sfu1, Sef1 and Hap43) and plays reciprocal roles in regulating *C. albicans* commensalism and pathogenesis (Chen et al., 2011). Specifically, the GATA family transcription factor Sfu1 was found to downregulate the expression of iron acquisition genes, prevent toxicity in iron-replete conditions, and contribute to intestinal commensalism of *C. albicans*. Sef1 acts as the central iron regulator for the expression of iron uptake genes in low-iron conditions and surprisingly, plays a dual role in regulating both intestinal commensalism and virulence (Chen et al., 2011). The CCAAT-binding repressor Hap43 transcriptionally represses Sfu1 which therefore de-represses iron acquisition gene expression in iron-limited conditions, and evidence has shown that mutant lacking *HAP43* exhibits a defect in virulence, supporting its role in pathogenicity (Chen et al., 2011; Hsu et al., 2011; Singh et al., 2011). Although the iron homeostasis regulatory circuit, as shown above, is essentially required for both commensalism and pathogenesis, the exact regulatory mode for each of the three factors and its application in driving the transition of *C. albicans* commensalism and pathogenicity, remains largely unsolved.

The CCAAT-binding factor is a highly conserved heteromeric transcription factor that specifically binds to the 5’-CCAAT-3’ consensus elements within the promoters of numerous eukaryotic genes (Kato, 2005). In mammals, three subunits, including NF-YA, NF-YB and NF-YC, form an evolutionarily conserved Nuclear Factor Y (NY-F) complex that exhibits the DNA-binding capacity to the CCAAT box and plays a vital role in transcriptional regulation of genes involved in proliferation and apoptosis, cancer and tumor, stress responses, growth and development (Dorn et al., 1987). A similar NF-Y-like (HAP) complex also exists in fungi like the budding yeast *Saccharomyces cerevisiae* and interestingly, this complex is composed of four subunits, termed Hap2, Hap3, Hap4 and Hap5 (Becker et al., 1991). Among them, the Hap2 (NF-YA-like), Hap3 (NF-YB-like) and Hap5 (NF-YC-like) subunits form a heterotrimeric complex that is competent for DNA binding, whereas the additional Hap4 subunit is an acidic protein and harbors the transcriptional activation domain necessary for transcriptional stimulation after interacting with the Hap2/Hap3/Hap5 complex (McNabb & Pinto, 2005). Interestingly, Hap4 is only present in fungi and functional analyses in a variety of fungal species identified that homologs of Hap4, such as the *Aspergillus nidulans* HapX, the *Schizosaccharomyces pombe* Php4, the *Cryptococcus neoformans* homolog HapX and *C. albicans* Hap43, were found to play both positive or negative roles in regulating the transcriptional responses to iron deprivation (Jung et al., 2010; Singh et al., 2011; Skrahina et al., 2017). Recently, there have been some progresses about the impact of Hap43 on the pathobiology of *C. albicans*. For example, Hap43 acts as a transcriptional repressor that is induced under low-iron conditions and required for iron-responsive transcriptional regulation and virulence, since knocking out *HAP43* in *C. albicans* significantly up-regulates the expression of genes involved in iron utilization under iron-limited conditions and attenuates virulence in a mouse model of disseminated candidiasis (Chen et al., 2011; Hsu et al., 2011). More importantly, genome-wide transcriptional profiling revealed that about 16% of the *C. albicans* ORFs were differentially regulated in a Hap43-dependent manner (Chen et al., 2011; Singh et al., 2011) and we found that a majority of differentially expressed genes (DEGs) are associated with oxidative stress and iron regulation, such as those involved in aerobic respiration, the respiratory electron transport chain, heme biosynthesis, and iron-sulfur cluster assembly, supporting the notion that Hap43 is one of the major iron-based redox sensors for *C. albicans* cells and contributes to the fine-tuned balance that adapts to different aspects of oxidative stress due to iron metabolism. Moreover, these data also raised a strong possibility that the regulatory function of Hap43 may be coupled to *C. albicans* commensalism by dealing with the cytotoxicity of ROS mainly generated in the iron-replete GI tract, in addition to its role in pathogenicity.

In this study, we sought to explore the underlying mechanism for a possible involvement of Hap43-dependent gene regulation in *C. albicans* gut commensalism, given that this commensal has to combat oxidative damage caused by excess iron content which is potentially detrimental for microbial cells. We unexpectedly unraveled an uncharacterized mechanism of posttranslational modification of the iron-responsive repressor Hap43 that regulates adaptation of *C. albicans* to commensalism in the gut by amelorating the iron-induced environmental oxidative stress.

## Results

### Deletion of *HAP43* significantly increases the commensal fitness of *C. albicans* in GI tract of mice fed a high-Fe diet

Accumulating evidence suggest the impact of the heterotrimeric CCAAT-binding complex on coordination of oxidative stress in fungi, as the HAP complex in *S. cerevisiae* activates the expression of genes involved in oxidative phosphorylation in response to growth on non-fermentable carbon source (Pinkham & Guarente, 1985) and the homologous complex (AnCF) in *A. nidulans* is regulated at the posttranslational level by the redox status of the cell and manipulates the expression of genes required for an appropriate response to oxidative stress (Thon et al., 2010). Moreover, microarray analyses in *C. albicans* showed that for 286 upregulated genes in *hap43*Δ/Δ relative to the wild type under iron-limiting conditions, 7.7% and 4.5% are those associated with aerobic respiration and the respiratory electron transport chain, respectively, highlighting the importance of Hap43 in iron-dependent oxidative stress (Chen et al., 2011). These observations prompt us to hypothesize that Hap43 may play an important role in regulating gastrointestinal commensalism of *C. albicans*, possibly by sensing changes of the oxidative status in this specific niche. To test this hypothesis, we first evaluated the contribution of Hap43 to the commensal fitness of *C. albicans* in GI tract, using a well-established mouse model of stable GI candidiasis (Chen et al., 2011). Groups of female C57BL/6 mice receiving a normal Fe diet (NFD) (37mg/kg Fe of diet) were inoculated by gavage with 1:1 mixtures of the wild type (WT) and *hap43Δ/Δ* mutant cells (***Figure. 1A***). The relative abundance of each strain in fecal pellets was monitored by qPCR using strain-specific primers (Table S2). Surprisingly, we did not observe significant differences in persistence between the WT and mutant (**Figure 1—figure supplement 1**). To investigate whether the inoculated *C. albicans* cells were really exposed to a host environment with high ROS levels, we examined ROS production in the colon tissue sections using the oxidant-sensitive fluorophore dihydroethidium (DHE). As shown in ***Figure. 1B and C***, the oxidative red fluorescence was almost undetectable in the colon (NFD), suggestive of insufficient ROS production. We therefore considered a possibility that the neglectable effect on ROS generation could be attributed to inadequate iron bioavailability in the gut. To test the possibility, we modified our animal model by changing the mouse diet from the normal chow to a high-Fe diet (HFD) (400mg/kg Fe of diet) (Mahalhal et al., 2018), since previous studies have shown that the amount of iron, which is about 10-fold higher than that in standard chow, is able to increase microbial exposure to iron without being overtly toxic to mice (Mahalhal et al., 2018; Schwartz et al., 2019). As expected, a three-day high-Fe diet (HFD) caused a significant increase of iron level in mouse colon, as determined by Prussian blue iron staining (***Figure. 1D***). In line with the elevated level of iron, we clearly observed a marked increase of ROS levels in mouse colon after a high-Fe diet, as detected by DHE showing an increase in fluorescence (***Figure. 1B and C***). The iron-induced ROS production in the gut was further confirmed by examining the transcript level of *DUOX2*. *DUOX2* encodes the dual oxidase 2, a hydrogen-peroxide generator at the apical membrane of gastrointestinal epithelia (Donko et al., 2014). qRT-PCR results showed that mice fed the high-Fe diet had significant increase in *DUOX2* mRNA levels in the colon compared with mice on a normal Fe diet (***Figure. 1E***). Finally, the hydrogen peroxide (H_2_O_2_) concentrations were determined in the mouse colon samples as an indicator of the level of oxidative stress (***Figure. 1F***). We observed that treatment of mice with HFD significantly increased levels of H_2_O_2_, indicating the induction of oxidative stress pathways in the colon tissues. Taken together, these data strongly support that a high-Fe diet is sufficient to sustain a persistent exposure of gut microbes to high levels of ROS.

**Figure 1.**
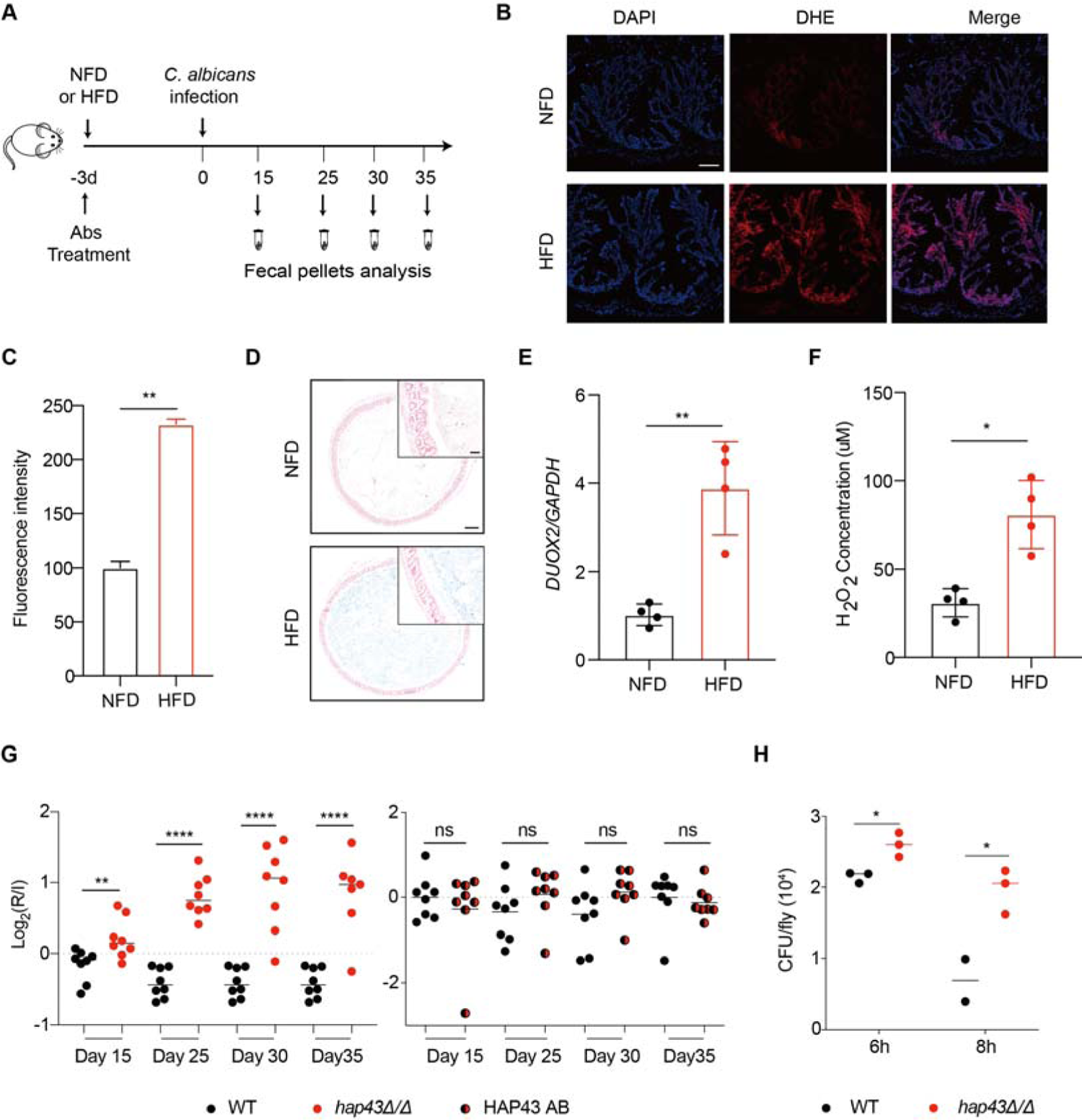
Deletion of *HAP43* significantly increases the commensal fitness of *C. albicans* in GI tract of mice fed a high-Fe diet. **(A)** As depicted in the schematics, mice were fed a normal Fe (NFD) or high-Fe diet (HFD) for 3 days prior to *C. albicans* inoculation. The mice continuously received the same diet during the course of experiments. **(B)** Colonic ROS accumulation in mice receiving a NFD or HFD diet for three days. Cryostat colonic sections were incubated with dihydroethidium (DHE) and DAPI. Scale bar, 100 μm. **(C)** Quantitative analysis using fluorescence intensity of DHE (a) in the colon. **(D)** Colonic samples were collected from mice fed either a NFD or HFD, formalin fixed, paraffin embedded, sectioned, and stained with Prussian blue for iron. Representative Prussian blue-stained colonic samples confirmed higher iron deposits in mice receiving HFD (iron blue, nucleus red). Scale bar, 200 μm; inset, 50 μm. **(E)** The expression of *DUOX2* mRNA in the colonic tissue of mice receiving NFD or HFD. Values were normalized to the expression levels of *GAPDH*. **(F)** The effect of iron on hydrogen peroxide (H_2_O_2_) levels in NFD or HFD-treated mice (n=4 mice per group). **(G)** Mutant lacking *HAP43* exhibits enhanced commensal fitness in HFD-treated mice. Mice (n=8 mice per group) fed a high Fe diet were inoculated by gavage with 1:1 mixtures of the wild-type (WT) and either *hap43Δ/Δ* mutant or *HAP43* reintegrant (*HAP43* AB) cells (1×10^8^ CFU per mice). The fitness value for each strain was calculated as the log_2_ ratio of its relative abundance in the recovered pool from the host (R) to the initial inoculum (I), and was determined by qPCR using strain-specific primers that could distinguish one from another. **(H)** Differences in fungal burden (expressed as CFUs) of flies assessed at different time points after incubation in a fresh vial containing live yeast media (4×10^8^ CFU of WT or *hap43Δ/Δ* mutant cells). Results from three independent experiments are shown. All data shown are means ± SD. ns, no significance; * *p*<0.05, \*\**p*<0.01, **** *p*<0.0001; by unpaired Student’s *t*-test (C, E, F, G) or two-way ANOVA with Sidak’s test (H). The following figure supplement for figure 1: **Figure 1 supplement 1.** Mutant lacking *HAP43* exhibits no change in commensal fitness in NFD-treated mice.

We repeated the competitive gut infections in mice receiving HFD, using the WT, *hap43Δ/Δ* mutant and *HAP43* reintegrant (*HAP43* AB) strains (***Figure. 1A***). Using this modified model, we found that a mutant lacking *HAP43* exhibited enhanced colonization fitness, such that the *hap43Δ/Δ* mutant cells significantly outcompeted WT *C. albicans* in 1:1 mixed infection (***Figure. 1G***), implying a negative impact of the iron-responsive regulator Hap43 in gut commensalism of *C. albicans*. This notion was further validated by using the wild-type *Drosophila melanogaster* as a model host to assess the gut fitness of the WT and *hap43Δ/Δ* mutant. Following a previously described protocol (Glittenberg et al., 2011), we set up *C. albicans* GI infections in the early third instar larvae of the common laboratory strain OrR of the WT or *hap43Δ/Δ* mutant. At time intervals of 6- and 8-hours post-infections, the infected flies were homogenized, serially diluted, and plated on plates for the recovery of fungal cells. Colony-forming unit (CFU) measurements indicated that, following 6 or 8 h of infection, the amounts of living mutant are much higher than WT *C. albicans* in the host (***Figure. 1H***). Taken together, our *in vivo* evidence highly suggests that Hap43 may play a negative role in regulating the gastrointestinal commensalism of *C. albicans,* especially under the circumstance in which the dietary stress is induced by ROS in the gut.

### High iron triggers Ssn3-mediated phosphorylation of Hap43

The microbial commensals colonizing the mammalian gut thrive on comparatively high levels of iron that are not digested and taken up by the upper intestine (Miret et al., 2003). Deletion of the iron-responsive regulator Hap43 results in a beneficial effect on *C. albicans* colonization in the gut, making it highly possible that iron influences the expression of Hap43. Indeed, we found that by both qRT-PCR and immunoblotting, Hap43 in WT strain was significantly less expressed in iron-repleted (H) medium in comparison to the iron-depleted (L) medium (***Figure. 2A and B***). Unexpectedly, immunoblot analysis of Hap43-Myc recovered from WT cells under iron replete vs depleted condition identified an increase in the electrophoretic mobility of Hap43 in iron-replete medium compared to that under iron-depleted conditions (***Figure. 2B***). Interestingly, when WT cells expressing Myc-tagged Hap43 were pre-grown to mid-exponential phase (OD_600_=0.4∼0.5) under iron-depleted conditions and then transferred into the iron-repleted medium (YPD), we found that a rapid gel mobility of Hap43 can be visualized at early time (2 mins) after medium change (***Figure. 2C***). We hypothesized that the shift in mobility on SDS-PAGE gel electrophoresis that is characteristic of the Hap43-Myc proteins might result from posttranslational modification, for example, a covalent phosphorylation event. To test this possibility, WT cells expressing Myc-tagged Hap43 were grown to mid-exponential phase in either iron-rich medium or iron-depleted medium, and cell lysates were treated with or without lambda phosphatase, a broad specificity enzyme which acts on phosphorylated serine, threonine and tyrosine residues. Immunoblotting analysis indicated that the mobility shifted form of Hap43-Myc reverted to the unshifted form if cell lysates were treated with lambda phosphatase, showing that the increased mobility induced by high iron was due to phosphorylation (***Figure. 2D***).

**Figure 2.**
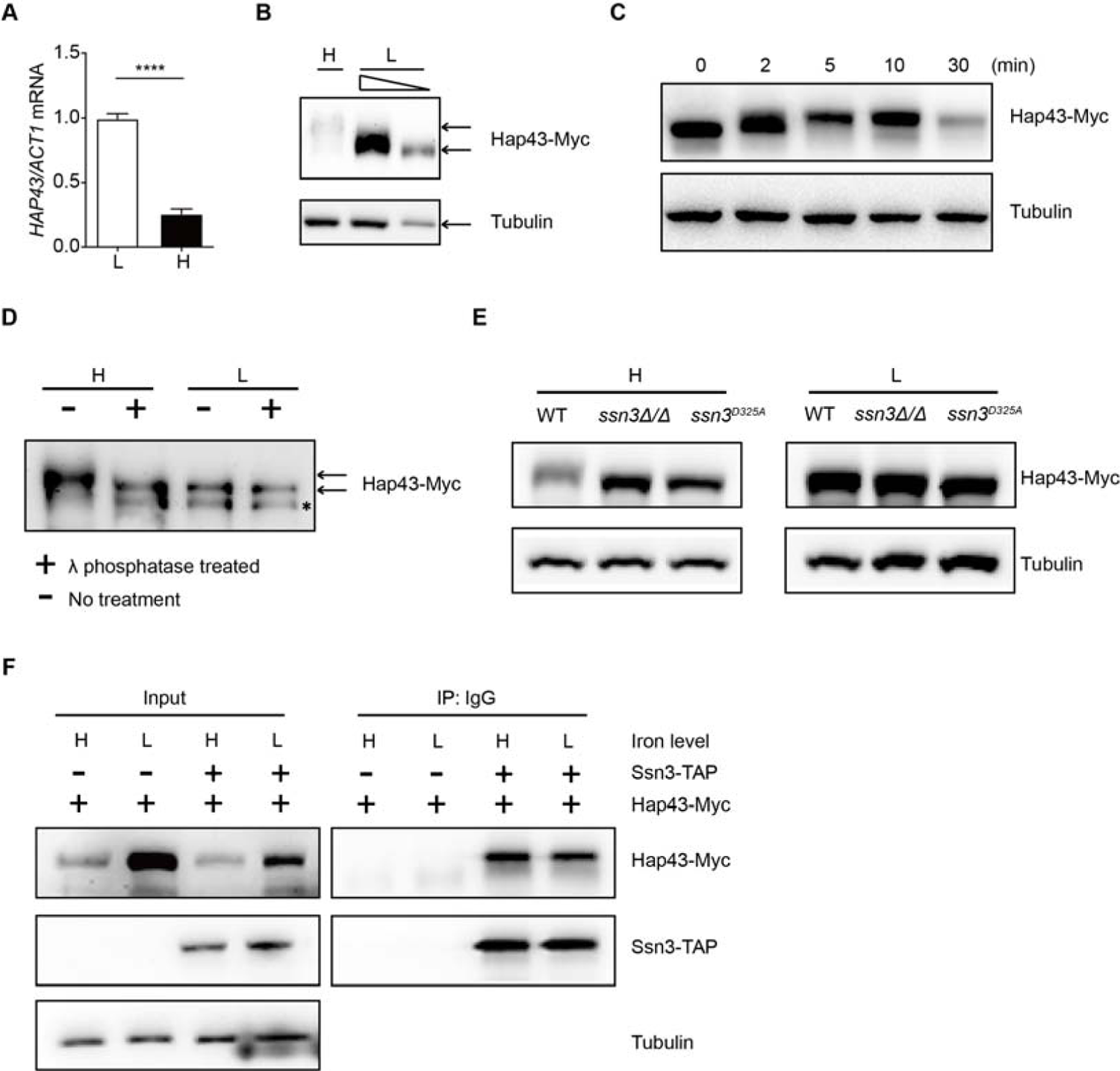
High iron triggers Ssn3-mediated phosphorylation of Hap43. **(A)** qRT-PCR analysis for *HAP43* mRNA in WT strain grown under iron-replete (H, high iron) or iron-depleted (L, low iron) conditions. Transcript levels were normalized to the level of *ACT1* mRNA. Results from three independent experiments are shown. All data shown are means ± SD. **** *p*<0.0001; by unpaired Student’s *t*-test. **(B)** Immunoblots of C-terminally tagged Hap43 (Hap43-Myc) in WT cells propagated under iron-replete (H) or iron-depleted (L) conditions. To better display the mobility-shift on protein, we added additional lane and loaded smaller quantities of total proteins from low-iron culture. α-tubulin, internal standard. **(C)** Time course for electrophoretic mobility of Hap43-Myc in WT cells during a shift from iron-depleted to iron-replete conditions. **(D)** Immunoblots of purified Hap43-Myc protein either treated (+) or not treated (-) with λ phosphatase. Note that higher amounts of total proteins from high-iron cultures were loaded. * indicates a presumed Hap43-Myc C-terminal proteolysis product. **(E)** Immunoblots of Hap43-Myc recovered from WT, *ssn3Δ/Δ* or *SSN3^D325A^* cells under iron-replete (H) or iron-depleted (L) conditions. α-tubulin, internal standard. **(F)** Hap43-Myc is co-immunoprecipitated with Ssn3-TAP. WT strains containing only Ssn3-TAP or both Ssn3-TAP and Hap43-Myc were grown under iron-replete (H) or iron-depleted (L) conditions. Lysates were prepared under nondenaturing conditions, and IgG-sepharose affinity column was used to immune-precipitate Ssn3-TAP and interacting proteins. The following source data for figure 2: **Figure 2-source data.** Uncropped images of gels and blots in Figure 2.

We previously reported that the Cys_6_Zn_2_ DNA binding protein Sef1, another key player operating in the iron-regulatory circuit of *C. albicans*, was phosphorylated under iron-depleted conditions and the phosphorylation was catalyzed by the protein kinase Ssn3 (Chen & Noble, 2012). To test whether Hap43 phosphorylation under iron-replete conditions also depends on the kinase activity of Ssn3, we expressed the Myc epitope-tagged version of Hap43 in *ssn3Δ/Δ* mutant strain and examined the mobility of Hap43 by immunoblotting.

Compared to that of WT, the higher mobility form of Hap43 under iron-replete conditions was abolished in the mutant lacking *SSN3*, supporting the role of Ssn3 in phosphorylation of Hap43 (***Figure. 2E***). An identical result was obtained when the mobility of Hap43-Myc was examined in the strain expressing a predicted kinase-dead allele of Ssn3 (Ssn3^D325A^) (***Figure. 2E***). Moreover, the putative enzyme-substrate interactions between Ssn3 and Hap43 was further reinforced through a co-immunoprecipitation assay, showing that Hap43-Myc was efficiently co-immunoprecipitated with Ssn3-TAP using either iron-replete or iron-depleted cells (***Figure. 2F***).

### Ssn3-mediated phosphorylation induces cytoplasmic localization and protein degradation of Hap43 by ubiquitin-proteasome pathway

Studies have shown that Ssn3 acts as a cyclin-dependent protein kinase and catalyzes the phosphorylation of a number of specific transcription factors that strongly contributes to their transcriptional activities, nuclear-cytoplasmic localization, and/or stability (Chi et al., 2001; Nelson et al., 2003). The effect of Ssn3-mediated phosphorylation on subcellular localization of Hap43 was investigated by indirect immunofluorescence. Under iron-depleted conditions, the localization of Hap43-Myc was primarily nuclear in both WT and *ssn3Δ/Δ* mutant strains (***Figure. 3A***). However, differences were observed under iron-replete conditions, in which the Hap43-Myc was found to be partially mislocalized from cytoplasm to nucleus in *ssn3Δ/Δ* mutant, compared to a complete cytoplasmic localization of this fusion protein in WT (***Figure. 3A***). The intracellular localization of Hap43-Myc in either WT or *ssn3Δ/Δ* mutant strains was further analyzed by immunoblot analysis of cell fractions. Yeast nuclei were purified using a modified method described previously (von Hagen & Michelsen, 2013) and the analysis of Hap43-Myc distribution showed that Hap43 is only detected in the nuclear fraction of *ssn3Δ/Δ* mutant cells but not WT, when cultures were grown under iron-replete conditions (***Figure. 3B***). These data highly suggested that blockade of Hap43 phosphorylation by *SSN3* deletion allows nuclear mislocalization of Hap43 in *C. albicans* grown under iron-replete conditions. In other words, Hap43 transcription factor is able to respond to iron status in *C. albicans* and modulates its expression and subcellular localization that depend on the posttranslational modification by covalent phosphorylation.

**Figure 3.**
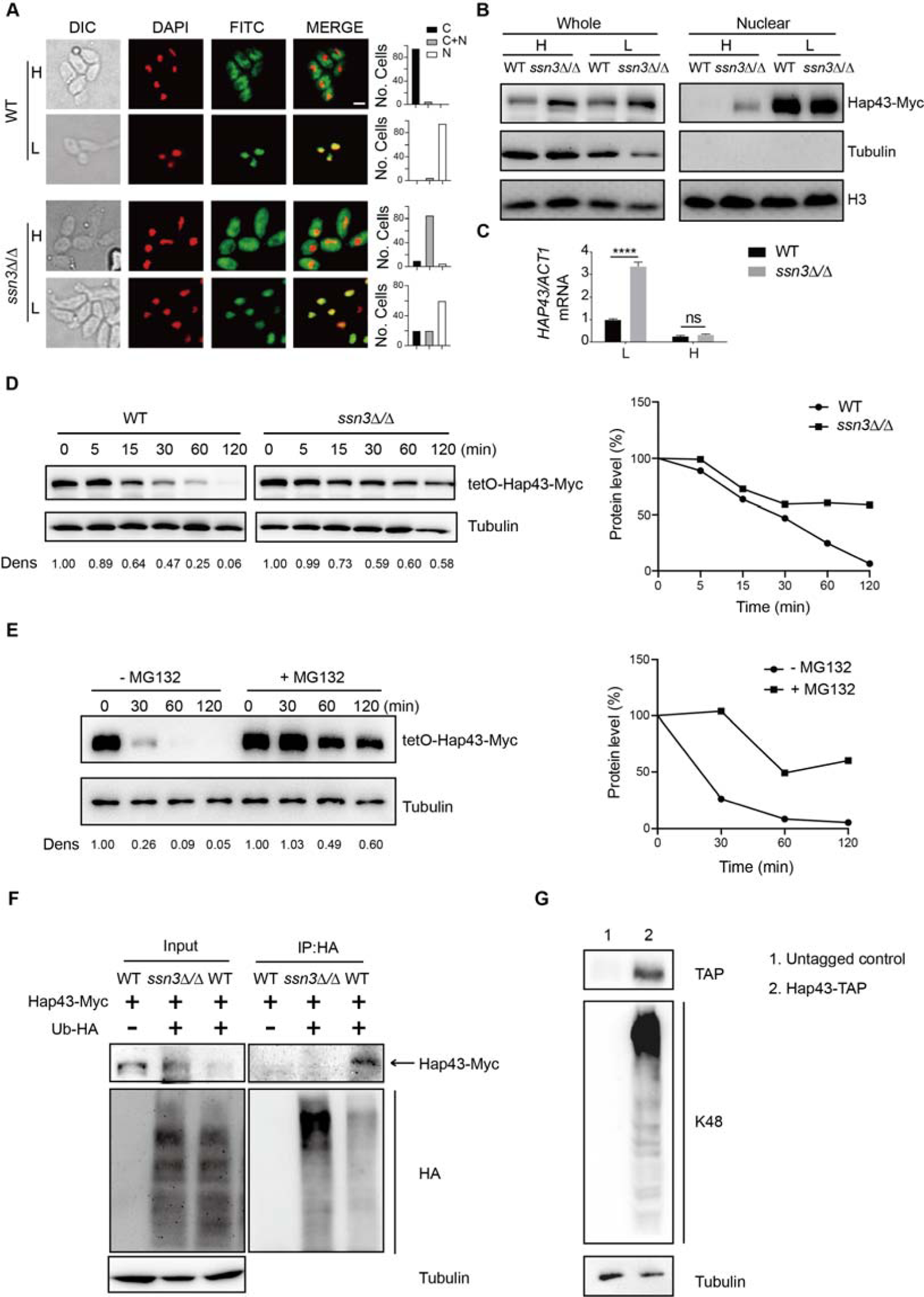
Ssn3-mediated phosphorylation induces cytoplasmic localization and protein degradation of Hap43 by ubiquitin-proteasome pathway. **(A)** Left panels: Indirect immunofluorescence of Hap43-Myc in WT and *ssn3Δ/Δ* mutant strains grown under iron-replete (H, high iron) or iron-depleted (L, low iron) conditions. DIC represents phase images, DAPI represents nuclear staining, FITC represents Hap43-Myc staining, and Merge represents the overlay of Hap43-Myc and nuclear staining. Right panels: Quantification of the cellular distribution of Hap43. Each bar represents the analysis of at least 100 cells. C representing >90% cytoplasmic staining, N >90% nuclear staining, and C+N a mixture of cytoplasmic and nuclear staining. Scale bar, 5 µm. **(B)** Immunoblots of Hap43-Myc in whole cell extracts and nuclear fraction of WT or *ssn3Δ/Δ* mutant cells propagated under iron-replete (H) or iron-depleted (L) conditions. Cellular contents were separated into cytosolic and nuclear fractions according to the protocol described in *Materials and Methods*. The nuclear marker H3 and cytoplasmic marker α-tubulin were used to display the purities of nucleus and cytoplasm. **(C)** qRT-PCR analysis for *HAP43* mRNA in WT and *ssn3Δ/Δ* strains grown under iron-replete (H) or iron-depleted (L) conditions. Transcript levels were normalized to the level of *ACT1* mRNA. Results from three independent experiments are shown. All data shown are means ± SD. ns, no significance; **** *p*<0.0001; by two-way ANOVA with Sidak’s test. **(D)** Hap43 protein is stabilized in a *ssn3Δ/Δ* mutant. WT or *ssn3Δ/Δ* strains stably expressing doxycycline-inducible Myc-tagged Hap43 (TetO-Hap43-Myc) were treated with doxycycline. Cells were harvested in the exponential phase of growth, washed to remove doxycycline, and resuspended in fresh iron-replete medium. The turnover of Hap43-Myc in WT or *ssn3Δ/Δ* cells was then evaluated following the tetO promoter shut-off by removal of doxycycline, through time-course experiments. Right panel: Hap43-Myc quantification after intensity analysis using Image J. **(E)** Similar to **D**, after treatment with doxycycline, WT cells (a copy of *ERG6* was deleted) stably expressing doxycycline-inducible Myc-tagged Hap43 (TetO-Hap43-Myc) were harvested, washed and treated with or without the proteasomal inhibitor MG132 (100 μM). The turnover of Hap43-Myc in WT cells was evaluated through time-course experiments. Right panel: Hap43-Myc quantification after intensity analysis using Image J. **(F)** Detection of Hap43 ubiquitination in *C. albicans*. WT and *ssn3Δ/Δ* mutant strains were engineered by stably expressing either Hap43-Myc alone or both Hap43-Myc plus tetO-HA-Ub. Both strains were incubated under iron-replete plus 50 μg/ml doxycycline conditions and cell extracts were subjected to immunoprecipitation with anti-HA-conjugated beads followed by Western blot analysis with anti-Myc antibodies for detection of ubiquitinated Hap43. **(G)** Detection of Hap43 polyubiquitination in *C. albicans*. The WT strain was engineered by stably expressing Hap43-TAP and grown under iron replete conditions. Cell extracts were immunoprecipitated with lgG-sepharose followed by Western blot analysis with anti-K48 linkage antibody for detection of K48-linked polyubiquitination of Hap43. The following source data and figure supplement(s) for figure 3: **Figure 3-source data.** Uncropped images of gels and blots in Figure 3**. Figure 3 supplement 1.** Chloroquine had no effect on Hap43 degradation under high iron conditions. **Figure 3 supplement 2.** Immunoblots showing the induction of an epitope-tagged 3xHA-ubiquitin under the control of the doxycycline (DOX) inducible promoter.

We note that loss of *SSN3* has a direct effect on the protein level of Hap43 when the cells were cultured in iron-replete conditions. Following the abolishment of increased mobility, the level of Hap43 is comparable with that found under iron-depleted conditions (***Figure. 2E***). Importantly, the increased steady level of Hap43 protein could not be explained by its transcriptional level, as we observed that deletion of *SSN3* has no effect on the mRNA level of *HAP43* under iron-replete conditions (***Figure. 3C***), suggesting that the posttranslational modification by covalent phosphorylation may promote protein instability of Hap43. To test this possibility, we employed the strains in which the sole Hap43-Myc allele is expressed under the control of the doxycycline (DOX) inducible promoter (TetO-Hap43-Myc/*hap43Δ*) in either WT or *ssn3Δ/Δ* mutant backgrounds. Exponential-phase cells growing in iron-replete (YPD) medium supplemented with 50 μg/ml doxycycline were harvested, washed and re-suspended in fresh YPD medium, and whole-cell protein extracts were prepared at each time point for analysis by Western blotting. Clearly, Hap43-Myc levels in WT were reduced by approximately 50% after 30 min incubation and continued to decline over the course of incubation (***Figure. 3D, left panel***). In comparison, abundance of Hap43-Myc in *ssn3Δ/Δ* mutant remained at a relatively high level during the treatment (***Figure. 3D, right panel***). These data highly suggested that phosphorylation of Hap43 mediated by Ssn3 kinase promotes its degradation.

In eukaryotic cells, lysosomal proteolysis and the ubiquitin-proteasome system represent two major protein degradation pathways mediating protein degradation (Lecker et al., 2006). To clarify the exact proteolytic pathway implicated in Hap43 turnover under iron-replete conditions, we incubated cells with specific and selective inhibitors of the lysosome (Chloroquine) or the proteasome (MG132). Previous studies have shown that proteasome inhibitors such as MG132 are unable to penetrate WT yeast cells due to the impermeability of the cell wall or membrane and therefore, mutant strains (e.g. *erg6Δ* and *pdr5Δ*) are required for experiments using the proteasome inhibitors since the mutant cells show increased drug permeability or reduced drug efflux (Tumusiime et al., 2011). We adapted the same strategy for inhibiting the proteasome and lysosome in *C. albicans*. A copy of *ERG6* gene was deleted from the strain in which the sole Myc-tagged version of Hap43 was expressed under the control of the doxycycline (DOX) inducible promoter (TetO-Hap43-Myc/*hap43Δ*), and the resulting mutant strain was treated with or without MG132. As shown in ***Figure. 3E***, treatment of mutant cells with 100 μM of MG132 for 30, 60, and 120 mins, significantly increased Hap43 protein levels compared with the untreated control. However, under the same experimental conditions, treating the cells with the lysosome inhibitor chloroquine had no effect on the decreased level of Hap43-Myc (**Figure 3—figure supplement 1**). Taken together, these data demonstrate that when *C. albicans* cells are grown under high iron conditions, the phosphorylated form of Hap43 is prone to be degraded through the proteasomal pathway.

To further verify this, we test a possibility of ubiquitination because this modification represents a common signal for proteasome-mediated protein degradation (Hershko & Ciechanover, 1998). In both WT and *ssn3Δ/Δ* mutant strain backgrounds (a copy of *HAP43* was C-terminally tagged with Myc epitope), we created strains that an epitope-tagged 3xHA-ubiquitin under the control of the doxycycline (DOX) inducible promoter was co-expressed with the Myc-tagged version of Hap43 (**Figure 3—figure supplement 2**). After a 6-h induction using doxycycline (50 μg/ml), log-phase cells were collected and lysed, followed by immunoprecipitation of whole cell extracts with anti-HA antibodies. Immunoblotting the precipitates with anti-Myc antibody revealed, as expected, a predominant band in WT but not *ssn3Δ/Δ* mutant (***Figure. 3F***), indicating that only the phosphorylated form of Hap43-Myc was able to bind ubiquitin. In the other direction, Hap43 was fused C-terminally to a tandem affinity purification (TAP) tag in the WT strain (Hap43-TAP/Hap43) and lysates were immunoprecipitated with IgG beads to recover the TAP-tagged Hap43 and the precipitates were immunoblotted with K48 linkage-specific polyubiquitin antibodies, considering the fact that the polyubiquitin chains linked through K48 are the principal signal for targeting substrates to the proteasome for degradation (Thrower et al., 2000). A reactive smear, characteristic of polyubiquitination, was associated with immunoprecipitated TAP-tagged Hap43 (***Figure. 3G***). Collectively, our data suggest that the Ssn3-mediated phosphorylation promotes protein degradation of Hap43 through a ubiquitin-proteasome pathway.

### Identification of the Hap43 phosphorylation sites that signal its ubiquitination and degradation

Together with the aforementioned results that the iron-responsive regulator Hap43 is phosphorylated in *C. albicans* cells grown under iron-replete conditions, this observation prompt us to identify at which serine/threonine residues Hap43 is phosphorylated. First, we started with an *in silico* prediction by using the Kinasephos 2.0 server (http://kinasephos2.mbc.nctu.tw/) (Wong et al., 2007) and this analysis predicted 12 putative serine/threonine phosphorylation sites within Hap43 of *C. albicans*. Moreover, Ssn3 of *C. albicans* is orthologous to *S. cerevisiae* Srb10, a cyclin-dependent kinase subunit of the Cdk8 module of Mediator (Bjorklund & Gustafsson, 2005), and putative Cdk8-dependent phosphorylation sites identified to date are serine/threonine residues flanked by a proline 1 to 2 residues toward the C-terminus, and/or by a proline 2 to 4 residues toward the N terminus (Chi et al., 2001). We examined the Hap43 sequence and identified another 17 potential Ssn3 kinase phosphorylation sites that meet the criteria described above (***Figure. 4A***). To experimentally confirm these in silico predictions, we generated amino acid substitution mutants in which the neutral amino acid alanine replaced serine/threonine at the predicted 12 putative phosphorylation sites to change the conserved phosphorylation motif in order to mimic the dephosphorylated state of Hap43. By use of the strain (TetO-Hap43-Myc/*hap43Δ*) described in ***Figure. 3D***, where the sole Hap43-Myc allele driven by the doxycycline (DOX) inducible promoter was expressed in the *hap43Δ/Δ* strain background, we successfully created seven mutants, each of which included one or two mutated S/T (to Ala) sites. In similar, exponential-phase cells growing in iron-replete (YPD) medium supplemented with 50 μg/ml doxycycline were harvested, washed and re-suspended in fresh YPD medium, and whole-cell protein extracts were prepared at indicated time points or after 2 h of incubation, and analyzed by Western blotting for the phosphorylation status and overall level of Hap43. Intriguingly, we found that single or double mutation of the predicted phosphorylation sites had no change of the phosphorylation pattern and consequently, still promoted protein degradation of Hap43 as the WT cells did (**Figure 4—figure supplement 1A and B**), indicating that phosphorylation of Hap43 should not occur in merely one or two residues.

**Figure 4.**
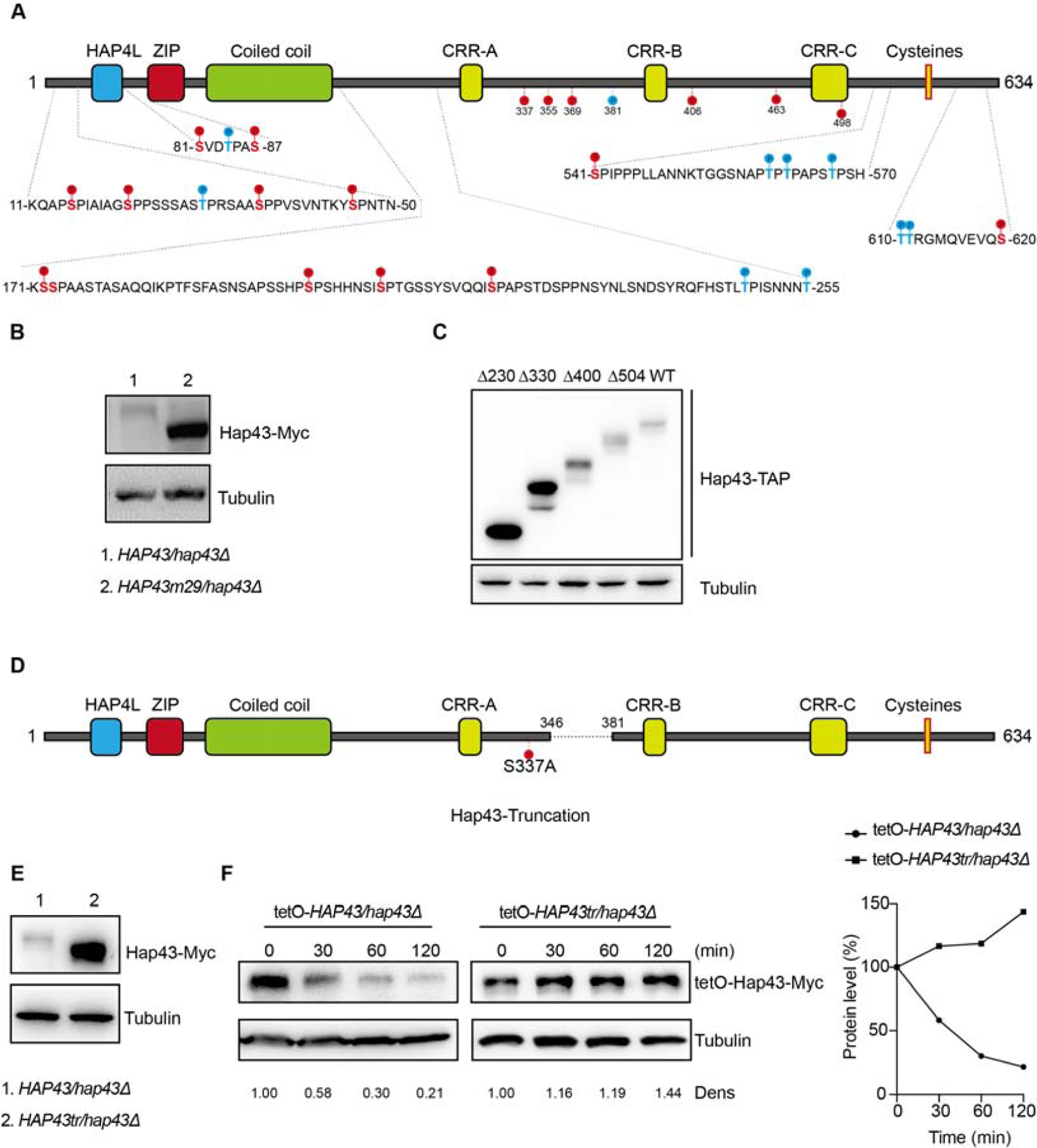
The critical phosphorylation sites are essential for Hap43 stabilization. **(A)** Schematic representation of *C. albicans* Hap43. Putative phosphorylation sites predicted by the Kinasephos 2.0 server and Cdk8-dependent phosphorylation sites are represented. **(B)** Immunoblots of Hap43-Myc in strains expressing either the WT or the amino acid mutation (*HAP43m*29; all 29 putative S/T phosphorylation sites were replaced with alanine residues) allele of Hap43. Cells were treated at high iron conditions. **(C)** Immunoblots of Hap43-TAP in WT and truncation mutant strains grown under iron-replete conditions. **(D)** Schematic representation of *C. albicans* Hap43 truncation. Hap43 truncation mutation (*HAP43tr*) was generated by deleting the 36 residues (346-381 aa) of Hap43 in HAP43^S337A^ strain. **(E)** Immunoblots of Hap43-Myc in strains expressing either the WT or the truncation mutation (*HAP43tr*) allele of Hap43. Cells were treated at high iron conditions. **(F)** Strains expressing either the WT or the truncation mutation (*HAP43tr*) allele of Hap43 under control of the inducible tetO promoter were treated with doxycycline. Cells were harvested in the exponential phase of growth, washed to remove doxycycline, and resuspended in fresh iron-replete medium (YPD). The turnover of Hap43-Myc in WT or truncation mutant cells was then evaluated following the tetO promoter shut-off by removal of doxycycline, through time-course experiments. Right panel: Hap43-Myc quantification after intensity analysis using Image J. The following source data and figure supplement(s) for figure 4: **Figure 4-source data.** Uncropped images of gels and blots in Figure 4**. Figure 4 supplement 1.** The Hap43 mutants harboring serine/threonine-to-alanine substitutions in its one or two putative phosphorylation sites showed the WT-like degradation patterns of Hap43 under high iron conditions. **Figure 4 supplement 2.** The mutants harboring amino acid substitutions or fragment truncation showed no defects in vegetative growth. Figure 4 **supplement 3.** The critical phosphorylation region is essential for Hap43 stabilization. **Figure 4 supplement 4.** The mutants harboring amino acid substitutions or fragment truncation showed no defects in vegetative growth. **Figure 4 supplement 5.** Identification of potential phosphorylation sites of Hap43.

We therefore generated a *HAP43* mutant strain (Hap43m-Myc/*hap43Δ*) that all 29 putative S/T phosphorylation sites, including the 12 residues predicted by computer algorithms and 17 residues matching the Cdk8 consensus phosphorylation sites, were replaced with alanine residues (***Figure. 4A***). An immunoblot with cell lysates from both WT (Hap43-Myc/*hap43Δ*) and *HAP43* mutant-29 (Hap43m29-Myc/*hap43Δ*) clearly revealed that the replacement of all 29 S/T residues abolished the upward shift (phosphorylation) of Hap43-Myc band induced by high iron and as a result, significantly increased the steady level of Hap43 (***Figure. 4B***). As a control, amino acid replacement appeared to have no effect on the growth and function of *HAP43* mutant harboring 29-point mutations under low iron conditions (**Figure 4—figure supplement 2**). Taken together, our experiments identified multiple S/T residues as important Hap43 phosphorylation sites *in vivo*.

Another alternative strategy for the role of phosphorylation is to assess the degradation of truncated Hap43. Four kinds of C-terminally deleted *HAP43* ORFs fused with the TAP tag were generated and introduced into *hap43Δ/Δ* mutant (**Figure 4—figure supplement 3A**). We showed that there was no significant difference in the transcript levels of WT and truncated *HAP43* (**Figure 4—figure supplement 3B**), however, their protein levels varied dramatically (***Figure. 4C***). Among them, Hap43 truncating mutations (Δ400 and Δ504) give rise to almost similar levels as the full length of WT, whereas the mutation (Δ330) results in a suddenly dramatic increase of Hap43 level. These results strongly suggest that the region within residues 330-400 harbors the signal contributing the phosphorylation-dependent degradation of Hap43. To further verify this, we deleted the 36 residues (346-381 aa) of Hap43 in HAP43^S337A^ strain (make sure there is no T or S left between 300-400aa) and generated a Hap43 truncation mutant (TetO-Hap43tr-Myc/*hap43Δ*) (***Figure. 4D***). Similar to the phenotypes observed in the *HAP43* mutant, deletion of the 36 amino acid residues also leads to increased level of the truncated form of Hap43 (***Figure. 4E***) and abrogated the high iron-induced protein degradation (***Figure. 4F***). As a control, fragment deletion appeared to have no effect on the growth and function of *HAP43* truncation mutant under low iron conditions (**Figure 4—figure supplement 4**).

By combining the results shown above, we finally focused on the four putative phosphorylation sites (S337/S355/S369/T381) between residue 336 and 381. To verify this, we generated a *HAP43* mutant-4 strain (Hap43m4-Myc/*hap43Δ*) in which site-directed mutagenesis was used to convert the codons for serine and tyrosine at these sites to condons for alanine (**Figure 4—figure supplement 5A**). Consistently, we found that under high iron conditions, the mutant exhibited significantly higher level of Hap43 proteins than that of the wild type (**Figure 4—figure supplement 5B**). As a control, replacement of these four residues with alanine appeared to have no effect on the growth and function of Hap43 (**Figure 4—figure supplement 5C**). Collectively, our data identified potential phosphorylation sites responsible for protein instability of Hap43 when *C. albicans* cells were grown under high iron conditions.

### Importance of Hap43 phosphorylation for alleviating Fenton reaction-induced ROS toxicity

Numerous studies have demonstrated that bivalent iron cation drives the Fenton reaction (Fe^2+^ + H_2_O_2_ Fe^3+^ + •OH + OH^-^) that plays an important role in the transformation of poorly reactive radicals into highly reactive ones, leading to many disturbances contributing to cellular toxicity (Ryan & Aust, 1992). The Hap complex, which is composed of Hap2, Hap3, Hap5 and Hap43 in *C. albicans*, has been found to play a key role in connecting the iron acquisition to oxidative stress response, by regulating the expression of oxidative stress genes (e.g., *CAT1*, *SOD4*, *GRX5* and *TRX1*), those who have been known to be induced in the production of ROS under iron-overloaded conditions (Mao & Chen, 2019). We therefore hypothesized that Hap43 phosphorylation may play a role in the coordinate regulation of *C. albicans* against iron-induced ROS toxicity. To test this, we first measured the intracellular ROS production in *C. albicans* cells grown under YPD or YPD supplemented with 200 μM FeCl_3_ conditions, by a fluorometric assay using hydroxyphenyl fluorescein (HPF; 5 μ) (Avci et al., 2016). As shown in ***Figure. 5A and B***, ROS levels were moderately elevated in *C. albicans* cells after incubation in YPD medium whereas massive increase was observed in medium supplemented with FeCl_3_, to a level comparable to that observed in medium with H_2_O_2_. As controls, iron-induced ROS production via Fenton reaction could be prevented by treating the cells with the antioxidant N-acetyl-L-cysteine (NAC). These data clearly indicated that high levels of iron are sufficient to significantly enhanced ROS production in *C. albicans*. More importantly, we observed that iron-triggered degradation of Hap43 could be inhibited by treating the cells with NAC (***Figure. 5C***) and treatment of *C. albicans* cells with menadione, an inducer of endogenous ROS, leads to the reduction of the Hap43 protein level (***Figure. 5D***), supporting the proposition that the promotion of iron-induced generation of ROS may account for the ubiquitin-dependent degradation of Hap43 after phosphorylation by Ssn3.

**Figure 5.**
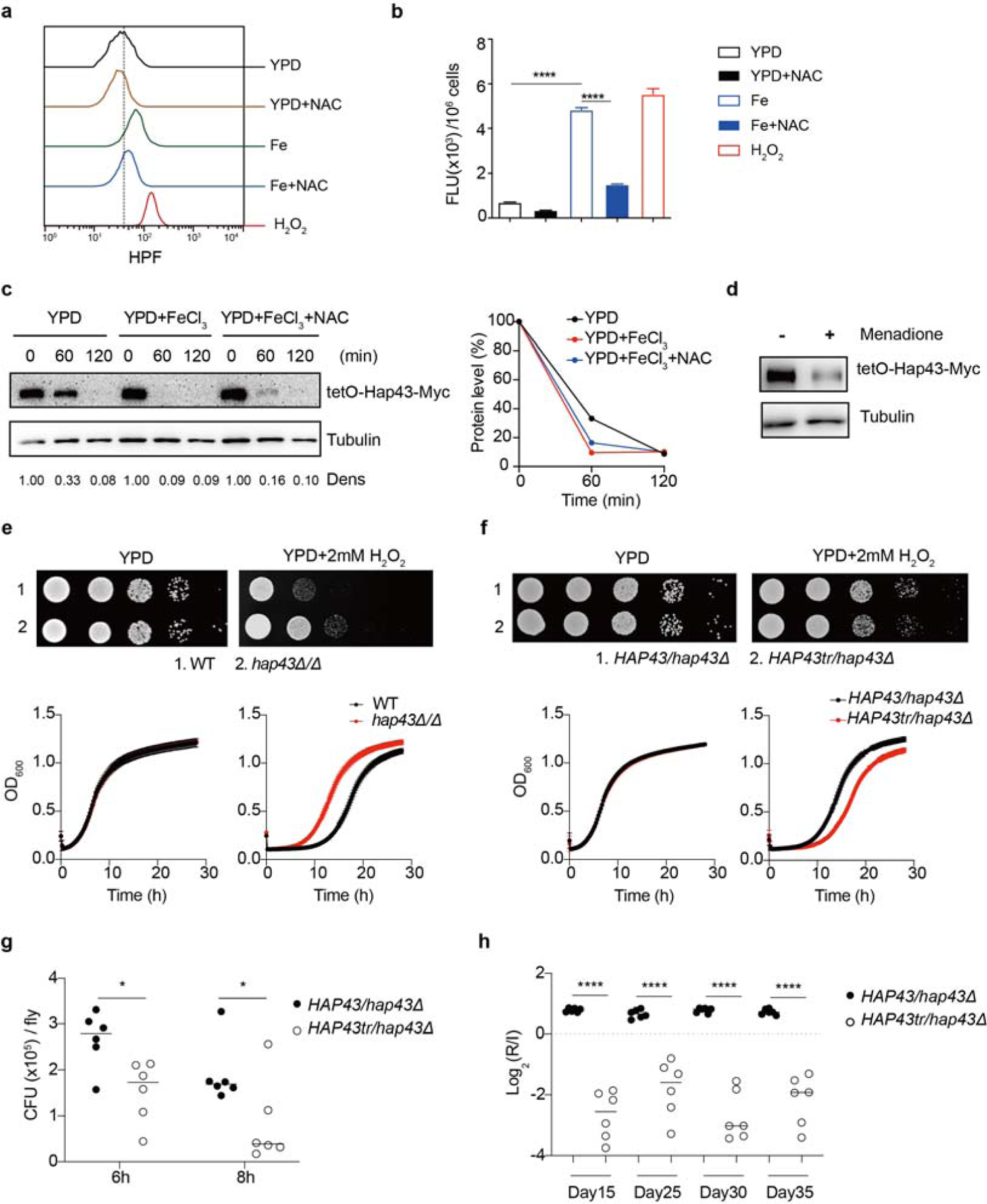
Hap43 phosphorylation is important for alleviating Fenton reaction-induced ROS toxicity and for GI colonization. **(A, B)** Intracellular ROS production of *C. albicans* under different experimental conditions. *C. albicans* yeast cells were grown on YPD supplemented with indicated reagents. About 1×10^7^ cells in exponential growth phase were collected, washed with PBS, stained with 5mM of HPF, and analyzed using FACS **(A)** or the microplate reader **(B)**. **(C, D)** Hap43 stability assay by immunoblots in WT strain stably expressing doxycycline-inducible Myc-tagged Hap43 (TetO-Hap43-Myc). Cells were incubated in YPD and YPD supplemented with 200 μM FeCl_3_, a combination of 200 μM FeCl_3_ and 20 mM N-acetyl-L-cysteine (NAC) (**C**) or 20 μM menadione for 120 min (**D**). **(E)** Growth of *hap43Δ/Δ* mutant under oxidative stresses. Top panel: WT and *hap43Δ/Δ* mutant cells were spotted with 10-fold serial dilutions onto YPD or YPD supplemented with 2 mM H_2_O_2_ and grown for 2 days at 30 ℃. Bottom panel: Growth curve analysis of WT and *hap43Δ/Δ* in YPD liquid medium supplemented with 2 mM H_2_O_2_ at 30 °C. OD_600_ readings were obtained every 15 min in a BioTek ^TM^ Synergy ^TM^ 2 Multi-mode Microplate Reader. **(F)** Growth of the truncation mutant (*Hap43tr*) under oxidative stresses. The experiments were conducted the same way as describe in **E**. **(G)** Similar to **Figure. 1G**, flies were fed on live yeast cells of indicated strains and the fungal burden of each strain (expressed as CFUs) was determined at different time points. **(H)** The truncation mutant of Hap43 exhibits decreased commensal fitness in mice. Similar to **Figure. 1F**, mice (n=6) were inoculated by gavage with 1:1 mixtures of the WT and the truncation mutant (*Hap43tr*) cells (1×10^8^ CFU per mice). The fitness value for each strain was calculated as the log_2_ ratio of its relative abundance in the recovered pool from the host (R) to the initial inoculum (I), and was determined by qPCR using strain-specific primers that could distinguish one from another. Results from three independent experiments are shown. All data shown are means ± SD. \**p*< 0.05; \*\*\*\**p*<0.0001; by one-way ANOVA with Sidak’s test **(B)**, two-way ANOVA with Sidak’s test **(G)** or unpaired Student’s *t*-test **(H)**. The following source data and figure supplement(s) for figure 5: **Figure 5-Source data.** Uncropped images of gels and blots in Figure 5**. Figure 5 supplement 1.** Hap43 phosphorylation is important for alleviating Fenton reaction-induced ROS toxicity and for GI colonization.

To further test the potential role of Hap43 phosphorylation in protecting *C. albicans* cells from ROS-induced cytotoxicity, we examined the growth of different strains (WT, *hap43Δ/Δ*, Hap43-Myc/*hap43Δ*, Hap43m29-Myc/*hap43Δ* and Hap43tr-Myc/*hap43Δ* mutants) in medium supplemented with or without H_2_O_2_. Compared to the WT, deletion of *HAP43* showed remarkable resistance to H_2_O_2_ (***Figure. 5E***), suggesting that loss of Hap43 promotes cell survival under oxidative stress. Actually, the observation is consistent with our *in vivo* fitness study showing an increased competitive ability of the *hap43Δ/Δ* mutant to colonize the GI tract (***Figure. 1***). Interestingly, we also found that compared to the WT (Hap43-Myc/*hap43Δ*), abolishment of Hap43 phosphorylation in each of the three mutants generated above, including the Hap43 truncation mutant (Hap43tr-Myc/*hap43Δ*), *HAP43* mutant-29 (Hap43m29-Myc/*hap43Δ*) or *HAP43* mutant-4 (Hap43m4-Myc/*hap43Δ*), showed significantly greater sensitivity to oxidative stress (***Figure. 5F***; **Figure 5—figure supplement 1A and B**), arguing that the ubiquitin-dependent degradation of Hap43 after phosphorylation contributes to the protection against ROS-induced cytotoxicity. This notion was further supported by the *in vivo* evidence that the *HAP43* mutant-29 and Hap43 truncation mutant could be outcompeted by the WT strain when cells stably colonize in both fly and mouse GI tract (**Figure 5—figure supplement 1C and D**; ***Figure. 5G and H***). Taken together, our data highly suggest that iron-induced Hap43 phosphorylation, followed by ubiquitin-dependent proteasomal degradation, acts to protect *C. albicans* from ROS toxicity and thus promote its survival in GI tract, a niche normally considered as an iron replete environment.

### Iron-induced phosphorylation and degradation of Hap43 leads to de-repression of antioxidant genes

ROS generation in actively growing cells occurs via Fenton reaction or as a byproduct of mitochondrial respiration. Previous studies have shown that Hap43 is primarily a transcriptional repressor and enriched in the nucleus in response to iron depletion, particularly responsible for repression of genes that encode iron-dependent proteins involved in mitochondrial respiration and iron-sulfur cluster assembly (Chen et al., 2011; Hsu et al., 2011). Moreover, our data revealed that iron-triggered posttranslational modification of Hap43, including cytoplasmic localization, phosphorylation, ubiquitination and proteasomal degradation, heavily impacts the ability of *C. albicans* to adapt and respond to oxidative stress. These findings are very informative and prompt us to examine whether the phosphorylation-dependent degradation of Hap43 may correlate with activation of antioxidant response. In other words, it is likely that iron-induced posttranslational modification of Hap43 may directly cause ROS elimination by upregulating the expression of antioxidant genes when *C. albicans* cells are bathed under conditions of high iron availability.

Given that the iron-responsive transcription factor Hap43 undergoes ubiquitin-dependent proteasomal degradation after phosphorylation, we provided evidence that deletion of the protein kinase Ssn3 prevents its degradation and causes nuclear mislocalization, when *C. albicans* cells are grown under iron-replete conditions (***Figure. 3A and B***). Consistently, replacement of all 29 S/T residues by alanine, as well as the truncated form, were found to abrogate phosphorylation and degradation of Hap43 (***Figure. 4***), prompting us to hypothesize that the unphosphorylated form of Hap43 through either amino acid substitutions or truncation, may alter its cellular localization when cells are grown under iron-replete conditions. To test this hypothesis, indirect immunofluorescence of formaldehyde-fixed yeast cells from WT (Hap43-Myc/*hap43Δ*), *HAP43* mutant-29 (Hap43m29-Myc/*hap43Δ*), or Hap43 truncation mutant (Hap43tr-Myc/*hap43Δ*) strain, at the early mid log phases of growth on YPD supplemented with FeCl_3_, was used to examine the subcellular localization of Hap43. As shown in ***Figure. 6A and* Figure 6—figure supplement 1A**, WT Hap43 localized to the cytoplasm, while unphosphorylated form of Hap43 (Hap43tr and Hap43m29) localized to the nucleus, suggesting that abolishing the phosphorylation-dependent modification resulted in relocation of Hap43 from cytoplasm to nucleus.

**Figure 6.**
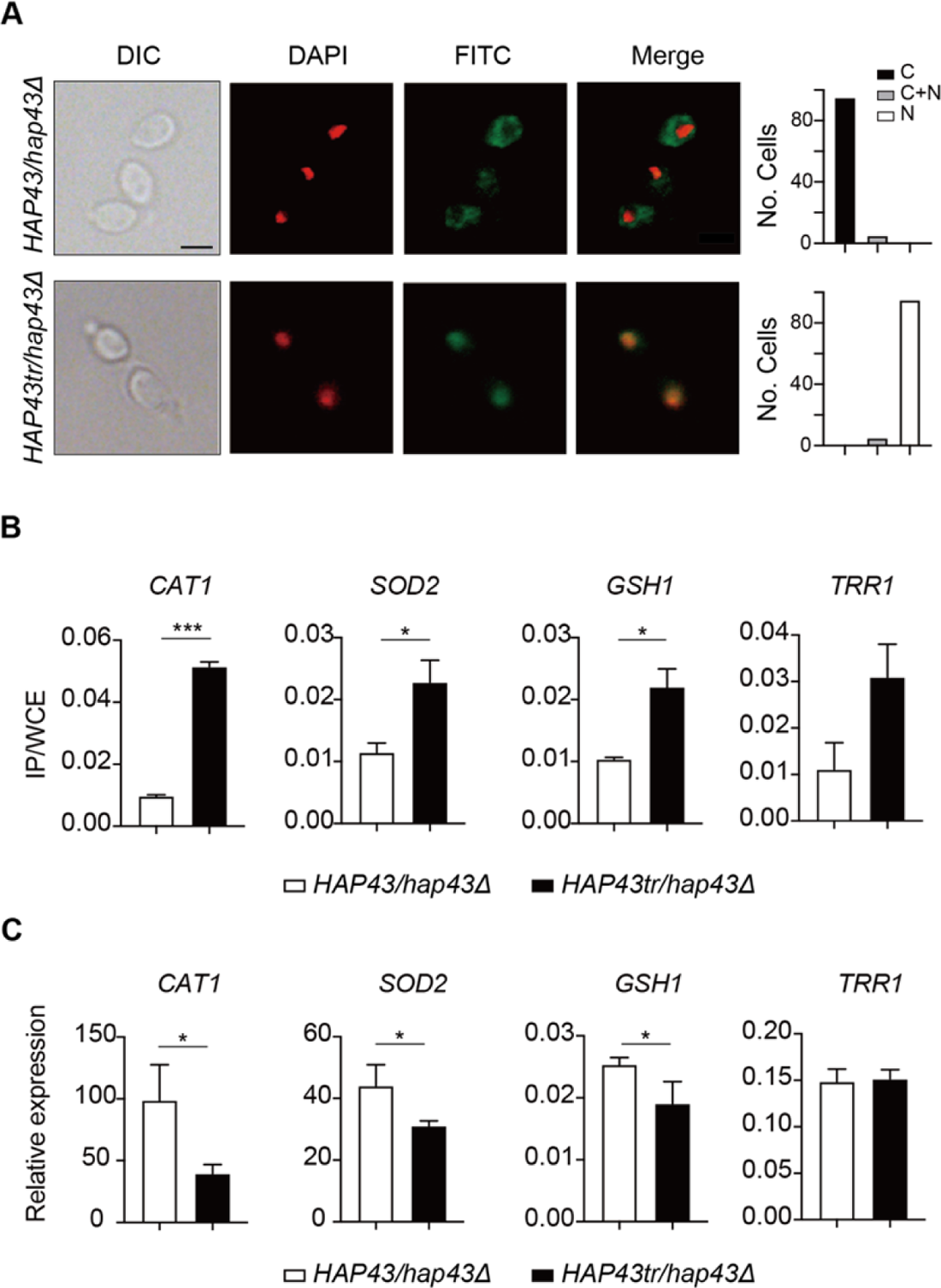
Iron-induced phosphorylation and degradation of Hap43 leads to de-repression of antioxidant genes. **(A)** Left panels: Indirect immunofluorescence of Hap43-Myc in *HAP43/hap43Δ* and *HAP43tr/hap43Δ* strains grown under iron-replete conditions. DIC represents phase images, DAPI represents nuclear staining, FITC represents Hap43-Myc staining, and Merge represents the overlay of Hap43-Myc and nuclear staining. Right panels: Quantification of the cellular distribution of Hap43. Each bar represents the analysis of at least 100 cells. C representing >90% cytoplasmic staining, N >90% nuclear staining, and C+N a mixture of cytoplasmic and nuclear staining. Scale bar, 5 µm. **(B)** ChIP of Hap43-Myc on the promoters that contain CCAAT boxes in a set of anti-oxidant genes. Overnight cultures of WT (*HAP43/hap43Δ*) and truncation mutant (*HAP43tr/hap43Δ*) cells were diluted in YPD plus 400 mM FeCl_3_ and grown to log phase at 30 ℃ before formaldehyde. Enrichment is presented as a ratio of qPCR of the indicated gene promoter IP (bound/input) over an *ACT1* control region IP (bound/input) of the tagged strain, further normalized to the control strain. **(C)** qRT-PCR analysis for mRNA levels of a set of antioxidant genes in WT (*HAP43/hap43Δ*) and truncation mutant (*HAP43tr/hap43Δ*) strains grown under iron-replete conditions. Transcript levels were normalized to the level of *ACT1* mRNA. Results from three independent experiments are shown. All data shown are means ± SD. ns, no significance; \**p*< 0.05; \*\**p*<0.01; \*\*\**p*<0.001; by unpaired Student’s *t*-test (B, C). The following figure supplement(s) for figure 6: **Figure 6 supplement 1.** Iron-induced phosphorylation and degradation of Hap43 leads to de-repression of antioxidant genes.

The antioxidant enzyme-mediated adaptive response has been demonstrated to attenuate toxicity caused by oxidative stress and a list of enzymes, including catalases, superoxide dismutases, peroxidases and peroxiredoxins, have been found to be the most ubiquitous effectors in microbial eukaryotes (Aguirre et al., 2005). Moreover, sequence analysis demonstrated the presence of CCAAT cis-acting element, a conserved Hap43 DNA recognition motif, on the promoter regions of antioxidant genes *CAT1*, *SOD2*, *GSH1* and *TRR1*. We therefore ask whether the unphosphorylated form of Hap43, once located to the nucleus, has the DNA binding capacity. ChIP-qPCR assays were performed to investigate Hap43 binding to the promoter sequences containing CCAAT motifs in the selected antioxidant genes. As expected, the mutated or truncated Hap43 (Hap43tr or Hap43m29) significantly enriched in the promoter regions of the four target antioxidant genes (***Figure. 6B and* Figure 6—figure supplement 1B**), which suggested a direct regulation of ROS detoxification in *C. albicans* by posttranslational modification of Hap43. Indeed, when the expression levels of these four antioxidant genes were examined by qPCR, we found that upon binding to the promoters directly, Hap43 significantly repressed the expression of *CAT1* and *SOD2* in both Hap43tr and Hap43m29 strains (***Figure. 6C and* Figure 6—figure supplement 1C**). Antioxidant enzymes such as superoxide dismutase (SOD) and catalase (CAT) form the first line of defense against ROS in organisms. Meanwhile, SOD is responsible for the formation of H_2_O_2_ through disproportionation to remove O_2_· and CAT metabolizes H_2_O_2_ into H_2_O and O_2_ (Ren et al., 2020; Van Breusegem et al., 2001). These results suggested that SOD and CAT may largely contribute to the resistance of *HAP43* mutants to iron-induced ROS.

Taken together, our data proposed a model (***Figure. 7; Graphical abstract***) that the iron-induced posttranslational modification of Hap43, including cytoplasmic localization, Ssn3-mediated phosphorylation, ubiquitination and proteasomal degradation, results in the de-repression of antioxidant genes (*e.g.*, *CAT1* and *SOD2*), an event that is most effective in lowering cytotoxicity induced by oxidative stress and promotes *C. albicans* commensalism in GI tract.

**Figure 7.**
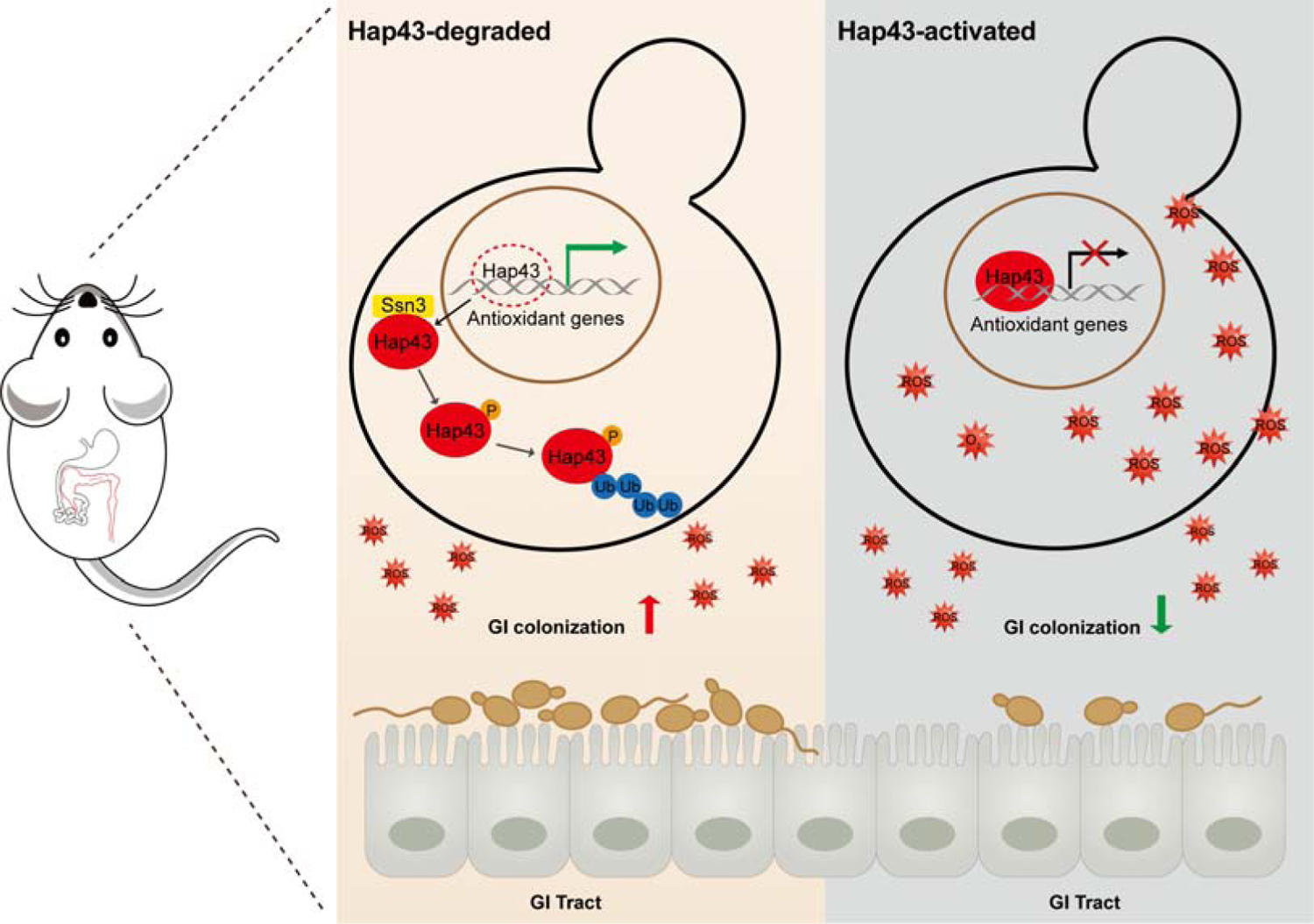
Model for the role of post-translational medication of Hap43 in promoting GI commensalism of *C. albicans*. In the iron rich environment such as GI tract, the iron-responsive regulator Hap43 is subject to covalent posttranslational modifications, including phosphorylation and ubiquitination, and causes cytoplasm-nuclear relocation and protein degradation via proteasome activity, thus serving as a positive signal to de-repress the expression of a set of antioxidant genes (*e.g.*, *CAT1* and *SOD2*), an event that is most effective in lowering cytotoxicity induced by iron-mediated ROS production and promotes *C. albicans* commensalism in GI tract.

## Discussion

Iron makes an ideal redox active cofactor for a variety of key biological processes and therefore becomes an indispensable element for all eukaryotes and the vast majority of prokaryotes. However, studies have revealed that iron excess is able to promote the production of potentially harmful ROS through accelerating the Fenton reactions, causing deleterious cellular effects such as lipid peroxidation, protein oxidation and carbonylation, and DNA mutagenesis and destabilization (Galaris et al., 2019). The need to avoid oxidative damages is particularly acute in the case of human fungal pathogens like *C. albicans*, mainly because these microbes are often subjected to assault by ROS produced by iron metabolism, environmental competitors or phagocytic cells during infections, as well as the endogenously produced ROS. Here, we discovered an uncharacterized detoxification strategy that *C. albicans* used to combat the toxic effects of ROS accumulation and promote its colonization in GI tract. Our data highly suggest that the iron-dependent global regulator Hap43, through a previously unknown posttranslational modification mechanism, senses the iron status of the cell and negatively regulates the gene expression of antioxidant enzymes.

Protein phosphorylation has been found to affect an estimated one-third of all proteins and recognized as the most widely studied posttranslational modification (Cohen, 2001). Changes in protein phosphorylation represent an important cell signaling mechanism that is frequently employed by cells to regulate the activities of transcription factors, for example, targeting for proteolytic degradation (Olsen et al., 2006). Moreover, a close connection between the ubiquitin-proteasome system and transcriptional activation has been reported in a number of studies (Lipford & Deshaies, 2003; Muratani & Tansey, 2003). Indeed, studies have shown that the posttranslational modification such as the phosphorylation-dependent ubiquitination and degradation is a highly conserved process across eukaryotes. For example, a powerful proteomic study in the budding yeast *S. cerevisiae* identified 466 proteins co-modified with ubiquitylation and phosphorylation (Swaney et al., 2013). A variety of extracellular stimuli in mammalian cells cause the rapid phosphorylation, ubiquitination, and ultimately proteolytic degradation of IκB, resulting in cytoplasm-nuclear translocation of NF-κB and induction of gene transcription (Ghosh & Dass, 2016). The same is the transcription factor SREBP1 who also undergoes phosphorylation and subsequent ubiquitination and degradation by the proteasome (Punga et al., 2006). In *Arabidopsis thaliana*, phosphorylation of the calmodulin-binding transcription activator 3 (CAMTA3) was found to trigger its destabilization and nuclear export (Jiang et al., 2020). Consistent with these observations, we described that under iron replete conditions, the iron-responsive transcription factor Hap43 of *C. albicans* undergoes ubiquitin/proteasome-mediated degradation upon a direct phosphorylation event mediated by Ssn3, a cyclin-dependent kinase previously known to have a similar activity on Ume6 degradation (Lu et al., 2019). Although the regions associated with phosphorylation and ubiquitination of Hap43 have been identified in our work, the precise modification sites remain unclear, more likely due to the presence of multiple modification sites and technical challenges such as the detection of low abundant proteins like transcription factors.

Previous studies have shown that ROS production in GI tract could be triggered by different abiotic or biotic stimuli. For example, iron (II) complex was found to interact with bile acids and the K vitamins to generate free radicals in the colon (Valko et al., 2001). The host’s defense through phagocytes induces an ROS burst that is required for pathogen killing and for regulating pro-inflammatory signaling in phagocytic cells (El-Benna et al., 2016). Moreover, similar studies have demonstrated that the antifungal action of different classes of antifungal compounds such as amphotericin B, miconazole, and caspofungin is related with the induction of ROS formation in fungi, especially in *Candida* species (Mello et al., 2011).

Previous studies showed that miconazole-mediated fungicidal activity against *C*. *albicans* was significantly inhibited by the addition of antioxidant (Kobayashi et al., 2002), and superoxide dismutase inhibitors enhanced the activity of miconazole against *C*. *albicans* biofilm cells (Bink et al., 2011). The ability of *C. albicans* to adapt and respond to oxidative stress is critical for its survival and virulence (Dantas Ada et al., 2015). Accumulating evidence have suggested that *C. albicans* cells respond to oxidative stress from the host environment through diverse strategies such as detoxifying ROS, repairing oxidative damages, synthesizing antioxidants and restoring redox homeostasis, and all of these actions involve the transcriptional induction of antioxidant genes encoding catalase (*CAT1*), superoxide dismutases (*SOD*), glutathione peroxidases (*GPX*) and components of the glutathione/glutaredoxin (*GSH1, TTR1*) and thioredoxin (*TSA1, TRX1, TRR1*) systems (Mao & Chen, 2019). Coincidently, we discovered the role of the transcription factor Hap43 in modulation of the transcription of antioxidant genes in response to iron. In iron replete environments (*e.g.*, host GI tract), Hap43 degradation leads to de-repression of antioxidant genes which enhances ROS detoxification and promotes the GI colonization of *C. albicans*.

Iron chelation has been explored as an adjunct in the treatment of fungal infections, particularly in salvage therapy (Reed et al., 2006). Some clinically approved iron-chelating drugs have been directly tested for inhibition of fungal pathogens, including *Cryptococcus*, *Rhizopu*, *Candida* and *Aspergillus* species (Symeonidis, 2009). For example, treatment of deferasirox, an FDA-approved iron chelator, significantly decreased the salivary iron levels and *C. albicans* CFUs of tongue tissue in a murine OPC model, and ultimately relieves neutrophil-mediated inflammation (Puri et al., 2019). Sepsis is a systemic inflammatory response induced by an infection (*e.g.*, bacteria or fungi), leading to organ dysfunction and mortality. During sepsis, iron homeostasis becomes disrupted and an excess of ROS is generated, causing damage to tissues. This can be potentially suppressed using iron chelators that selectively bind iron to prevent its participation in ROS-associated inflammatory reactions. Given the importance of Hap43 degradation in ROS detoxification and *C. albicans* commensalism, it is plausible that iron chelator therapy by blocking the process of protein phosphorylation and degradation could be considered as an alternative therapeutic approach against invasive fungal infection. Our findings may deliver new clues for the development of innovative drugs to fight invasive fungal infection.

## Materials and Methods

### Ethics statement

All animal experiments were carried out in strict accordance with the regulations in the Guide for the Care and Use of Laboratory Animals issued by the Ministry of Science and Technology of the People’s Republic of China. All efforts were made to minimize suffering. The protocol was approved by IACUC at the Institut Pasteur of Shanghai, Chinese Academy of Sciences (Permit Number: A160291).

### Media

*C. albicans* strains were grown at 30 °C in YPD (1% yeast extract, 2% Bacto peptone, 2% glucose) or SD (0.67% yeast nitrogen base plus 2% dextrose) as ‘iron-replete’ medium. ‘Iron-depleted’ medium is YPD or SD supplemented with one of the specific iron chelators, 500 μM bathophenanthroline disulfonic acid (BPS). Doxycycline (50 μg/ml) was added to YPD for Tet-induced expression. When required, MG132 (100 μM) or Chloroquine (100 mM) was added to growth medium to inhibit protein degradation.

### Plasmid and strain construction

SC5314 genomic DNA was used as the template for all PCR amplifications of *C. albicans* genes. The *C. albicans* strains used in this study are listed in Table S1A. The primers used for PCR amplification are listed in Table S2. Plasmids used for Hap43-Myc tagging and knockout gene complementation are listed in Table S1B. Construction of *C. albicans* knockout mutants, complemented strains, strains expressing Myc-tagged Hap43 fusion protein, and overexpression strain for Hap43 was performed as previously described (Chen et al., 2011).

To generate tetO-Hap43-Myc strains (CB247), we used the pNIM1 and replaced the caGFP reporter gene by *HAP43*-13xMyc. *HAP43*-13xMyc was amplified with primers (CBO838 and CBO839) that introduced SalI and BglII sites from pSN161. The PCR product was appropriately digested and inserted into SalI/BglII-digested vector pNIM1 to generate pCB127. *hap43Δ/Δ* strain (SN694) was transformed with the following gel-purified, linear SacII-KpnI digested DNA fragments from pCB127. To generate tetO-HA-Ub strain (CB453 and CB494), 3xHA-Ubiquitin (*Saccharomyces cerevisiae*) was synthesized by company (Gen Script Nanjing Co.,Ltd.). The plasmid was appropriately digested and inserted into SalI/BglII-digested vector pNIM1 to generate pCB193. Hap43-Myc strain (SN856) or Hap43-Myc, *ssn3Δ/Δ* strain (CB12) was transformed with the following gel-purified, linear the SacII-KpnI digested DNA fragments from pCB193 respectively.

### In vitro growth assay

For agar plate assays, fresh overnight yeast cultures were washed and diluted in PBS to adjust the optical density (OD_600_) to 1.0. Then 10-fold serial dilutions were prepared and 5 μl aliquots of each dilution were spotted onto appropriate agar plates. For growth curves in liquid media, cells from overnight cultures were diluted to a starting OD_600_ of 0.15 into the indicated medium. At indicated time intervals optical densitiy at 600 nm (OD_600_) was measured. Presented data (means and SDs) from three technical replicates were shown and plotted in Graphpad Prism.

### Fluorescence microscopy

*C. albicans* was grown at 30 ℃ for 5∼6 hours in YPD or YPD supplemented with 500 μM BPS medium to OD600 = 0.8∼1.0. Cells were fixed by 4.5% formaldehyde for 1 hour and digested by 80 μg/ml Zymolase-20T in 37 ℃ for 15 min. Cells were transformed to polylysine-d coated culture dishes and remove most of the un-attached cells. To flatten cells, add pre-cold (−20 ℃) methanol for 5 min followed by pre-cold (−20 ℃) acetone for 30s. The 9E10 anti-c-Myc antibody was used at a 1:150 dilution overnight after cells were completely dry. A 1:400 dilution of Cy2-conjugated secondary antibody was used for 1h. Images were acquired under oil objective using an inverted microscope. DIC, DAPI and FITC images acquired.

### Promoter shutdown assays

*C. albicans* strains containing Hap43-Myc or 3xHA-Ubiquitin under the regulation of the tetO promoter were grown in YPD at 30 °C overnight, then diluted 1:100 into YPD plus 50 μg/ml doxycycline to induce the expression of Hap43-Myc. Then the medium was replaced by fresh YPD medium at 30 °C to shut off the promoter. Aliquots were collected after the times indicated.

### Protein extraction and immunoblotting

*C. albicans* protein extracts were prepared under denaturing conditions. Briefly, lysates corresponding to 1 OD_600_ of cells were analyzed by SDS-PAGE and immunoblotted with either anti-c-Myc (9E10, Covance Research) for Myc-tagged proteins or anti-peroxidase soluble complex antibody (Sigma, P2416) for TAP-tagged proteins. Immunoblots were also probed with anti-alpha tubulin antibody (Novus Biologicals, NB100-1639) as a loading control. At least three biological replicates were obtained for each experiment shown and ImageJ software was used for densitometry.

### Lambda phosphatase treatment

100 ml *C. albicans* cells in log phage was washed with 1ml ice-cold 1.2M sorbitol twice and split into two tubes. Add 500 μl protein extraction buffer (420 mM NaCl, 200 μM EDTA, 1.5 mM MgCl_2_, 10% Glycerol, 0.05% Tween-20, 50 mM Tris-Cl, pH 7.5) containing 0.5M fresh-made DTT and protease inhibitor cocktails (Roche, USA). Add 0.5mm glass beads and break cells by vortex (6x 30s, 2 min interval on ice, top speed, 4 °C). Spin at top speed and transfer supernatant to a new tube. Add 5 μl 10x PMP buffer, 5 μl 10 mM MnCl_2_ buffer and 0 or 2 μl lambda phosphatase (NEB #P0753S, USA) in 38 μl supernatant. The mix was incubated for 60 min at 30 °C, followed by 10 min in 65 °C.

### Immunoprecipitation and pull-down assay

100 ml cells expressing TAP-tagged Hap43 or Ssn3 as well as cells expressing HA-tagged ubiquitin were collected by centrifugation in log phage. Cells were washed three times with ice-old water, and resuspended in 1 ml of lysis buffer (20 mM Tris, pH 7.4, 100 mM KCl, 5 mM MgCl_2_, 20% glycerol) with protease and phosphatase inhibitors (Roche). Cells were lysed using a Bead Beater and 300μl of glass beads. Cell lysates were centrifuged for at max speed at 4 °C for 15min. Protein concentration of the supernatants was measured by the Bradford assay and whole cell extracts were collected in freezer. 3 mg of proteins was used for immunoprecipitation with 50 ml of immunoglobulin G-Sepharose resin (IgG Sepharose 6 Fast Flow, GE Healthcare) or Anti-HA affinity matrix beads (Roche, USA). After protein overnight rotation at 4°C, the resin was washed 3 times with lysis buffer. For TAP-tagged proteins, the resin was washed twice with tobacco etch virus (TEV) protease cleavage buffer (10 mM Tris-HCl, pH 8, 150 mM NaCl, 0.5 mM EDTA, 0.1% Tween-20). Halo TEV protease (Promega, USA) cleavage was performed in 1 ml buffer at 4 °C overnight. The TEV eluate was collected and proteins were recovered by TCA (trichloroacetic acid) precipitation. For HA-tagged proteins, the resin was boiled in SDS-PAGE loading buffer (50 mM Tris-HCl, pH 6.8, 2% SDS, 10% (v/v) Glycerol, 2 mM DTT, 0.01% (w/v) Bromophenol Blue).

### Nuclear fraction separation

The nuclear fraction was prepared as a described method (von Hagen & Michelsen, 2013). 50 ml *C. albicans* cells in log phage was washed with preincubation buffer (100 mM PIPES-KOH pH 9.4, 10 mM DTT). The pellet was resuspended preincubation buffer and incubate in 30 °C for 10 min. Spin down and resuspend in 2 ml Lysis buffer (50 mM Tris-HCl pH 7.5, 10 mM MgCl_2_, 1.2 M sorbitol, 1 mM DTT) plus 80 μl Zymolase 20T (2.5 mg/ml) and incubate in 30 °C for 60 min. Cells were centrifuged and washes in lysis buffer twice and were resuspended in 2 ml Ficoll buffer (18% Ficoll 400, 100 mM Tris-HCl, pH 7.5, 20 mM KCl, 5 mM MgCl_2_, 3 mM DTT, 1 mM EDTA) containing protease inhibitors as described above. The cells were lysed using a Dounce homogenizer. Unlysed cells and cell debris were removed by 3000 rpm 15 min centrifugation. The supernatant was equally divided in two portions: one was saved as whole cell extract and the other was centrifuged at max speed for 15 min. The pellet (nuclear) was washed by PBS twice and resuspended in SDS loading buffer.

### Fungal genomic DNA isolation and total RNA preparation for RT-qPCR

Fungal genomic DNA isolation was performed as previously described (Chen & Noble, 2012). Samples were harvested by centrifugation and the pellets were resuspended in a DNA extraction solution containing 200 μl of breaking buffer (2% Triton X-100, 1% SDS, 100 mM NaCl, 10 mM Tris-HCl pH 8.0, 1 mM EDTA pH 8.0), 200 μl of acid Phenol: Chloroform: Isoamyl alcohol (pH 8.0, Ambion) and a slurry of acid-washed glass beads (Sigma-Aldrich). Fungal cells were mechanically disrupted with a FastPrep-24™ 5G (MP Biomedicals, USA) and genomic DNA were extracted and precipitated with isopropanol.

Fungal RNA was prepared as described previously (Chen & Noble, 2012), 1–2 μg of each RNA was treated with RNase-free Dnase I (Promega, Madison WI, USA) and reverse transcribed using the PrimeScript RT reagent Kit (TaKaRa). qPCR was performed using the SYBR Green Master Mix (High ROX Premixed) (Vazyme, Nanjing, China) using the primers in Table S2. Normalization of expression levels was carried out using the *ACT1* genes and the primers for *ACT1* was used as previously described. At least three biological replicates were performed per strain per condition.

### Chromatin Immunoprecipitation

ChIP experiments were performed essentially as described (Nobile et al., 2009). Unless otherwise noted, cells were crosslinked with 1% formaldehyde for 20 min at 30 ℃, followed by 125 mM glycine for 5 min. Cell pellets were resuspended in 700 ul ice-cold lysis buffer (50 mM HEPES-KOH pH 7.5, 140 mM NaCl, 1 mM EDTA, 1% Triton X 100, 0.1% NaDOC) with protease inhibitor cocktails (Roche, USA). Vortex with 300 ul glass beads at max speed for ∼2hr at 4 ℃. Recover the lysate by inverting and centrifuging the tubes with punctures on bottom and top of tubes by a 26G needle. Hap43-Myc were immunoprecipitated with 2–5 mg antibody (anti-Myc, 9E10, Covance Research) from lysates corresponding to optical density 600 (OD_600_) of cells at 4 ℃ overnight. Add 50 ul of prepared A or G beads suspension to each IP sample and rotate for 2 hr at 4 ℃. Wash twice with lysis buffer, high salt lysis buffer (50 mM HEPES-KOH pH 7.5, 500 mM NaCl, 1 mM EDTA, 1% Triton X 100, 0.1% NaDOC) and wash buffer (10 mM Tris-Cl pH 8.0, 250 mM LiCl, 0.5% NP-40, 0.5% NaDOC, 1 mM EDTA) respectively and resuspend in TE buffer. Products were eluted in elution buffer and incubated in 65 ℃. DNA was de-crosslinked by proteinase K (Sigma, USA) and 4 M LiCl and purified by phenol: chloroform: isoamyl alcohol. Immunoprecipitated DNA was quantified by real-time PCR (qPCR) with primers and normalized against *ACT1*.

### Measurement of ROS production

3’(4-Hydroxyphenyl)-fluorescein (HPF; Molecular Probes, OR, USA) was used for detecting •OH production (Avci et al., 2016). Log-phased *C. albicans* cells were washed in PBS buffer twice. HPF fluorescent probe was added to washed cells and kept in a 37 ℃ shaker for 30 min. Subsequently, cells were centrifuged (3200 rpm, 3 min) immediately and were resuspended in PBS. The stained cells were detected by a fluorescent microplate reader (Thermo Fisher, USA) or BD LSR Fortessa flow cytometer (BD Bioscience). Cells were also counted using a hemocytometer. The relative fluorescence density of each sample was calculated as FLU divided by the cell number to evaluate intracellular ROS levels.

### Determination of Colonic H_2_O_2_

Determination of H_2_O_2_ was performed according to the protocol of Beyotime kit (Cat #S0038, Beyotime, China). Briefly, 50 mg of colon tissue fragment was homogenized with 200 ul of lysis buffer and was centrifuged at max speed for 5 min in 4 ℃. Subsequently, 50 ul of supernatant was mixed with 100 ul test buffer and incubate for 30 min in room temperature. Then A560 was detected by a fluorescent microplate reader (Thermo Fisher, USA). Readings were calculated by the standard curve that was prepared from three series of calibration experiments with 5 increasing H_2_O_2_ concentrations (range 1-100 uM/l).

### *Drosophila* infection assays

The gut infection assays were performed as described previously (He et al., 2017). The oreR flies were used as our wild-type background flies. All flies were maintained on maize malt molasses food in bottles and reared pre-infection at 25 °C under a normal light/dark cycle. Mid-log phase *C. albicans* cells were harvested and resuspended in 5% sucrose and adjusted to an OD_600_ of 100. 3-5 days old male flies were dehydrated for 2 h without food and water, and then transferred into a vial covered with filter paper soaked with 5% sucrose solution containing the indicated *C. albicans* cells. Flies that had fed on 5% sucrose only were used as control. Both infected and control flies were incubated at 29 °C and transferred to conventional food after infection. Flies were ground in an Eppendorf tube with 200 μl of PBS 6 or 8 hours after infection with pipette tips and serial dilutions of the homogenates were plated onto YPD agar for CFUs.

### Mice infection assays

Female C57BL/6 mice (6-8 weeks old, weighing 18-20g) were purchased from Beijing Vital River Laboratory Animal Technology Company (Beijing, China). The mice were routinely maintained in a pathogen-free animal facility at Institut Pasteur of Shanghai, Chinese Academy of Sciences. All mice had free access to food and water in a specific pathogen-free animal facility with controlled temperature, humidity and a pre-set dark-light cycle (12 h: 12 h). Infections were performed under SPF conditions. The “normal diet” was a standard chow diet (37mg/kg iron; Shanghai SLAC Laboratory Animal Co.,Ltd). The “high-iron diet” was a diet supplemented with 400mg/kg iron (Shanghai SLAC Laboratory Animal Co.,Ltd). For competed infection, mice were received penicillin (1.5 mg/ml) and streptomycin (2 mg/ml) in their drinking water for 3 days prior to gavage with 1 × 10^8^ CFUs of 1:1 mixtures. Stool samples were homogenized in PBS and cultured in Sabouraud plates supplemented with ampicillin 50 μg/ml and gentamicin 15 μg/ml. Genome DNA were was extracted for fitness value of each strain by qPCR using strain-specific primers. For iron staining, colons were fixed with 10% formalin, and paraffin-embedded sections were stained with fresh-made Prussian blue staining solution. For ROS staining, colons were ‘snap-frozen’ in optimum cutting temperature compound, and frozen sections were stained with dihydroethidium (DHE) and DAPI. For colonic RNA, RNA was extracted with TRIzol according to the manufacturer’s instructions (Invitrogen).

### Statistical analysis

Data were presented as mean ± SD for continuous variables. All statistical analyses were performed with GraphPad Prism 8 (San Diego, Calif, USA) and details were provided in the Figure legends. The following *p*-values were considered: \**p* < 0.05; \*\**p* <0.01; \*\*\**p* < 0.001; \*\*\*\**p* < 0.0001.

## Acknowledgements

C.C. is supported by grants from the MOST Key R&D Program of China (2022YFC2303200; 2022YFC2304700); the National Natural Science Foundation (32170195); the Shanghai Municipal Science and Technology Major Project (2019SHZDZX02); the Open Project of the State Key Laboratory of Trauma, Burns and Combined Injury, Third Military Medical University (SKLKF201803); the Innovation Capacity Building Project of Jiangsu Province (BM2020019). H.L. is supported by the Special Program of PLA (19SWAQ18). Y.W. is supported by a National Postdoctoral fellowship (No.312780) and Shanghai Post-doctoral Excellence Program (No.2022647). X.H. is supported by grants from the National Natural Science Foundation (32070146, 31600119); the Natural Science Foundation of Shanghai (20ZR1463800, 15ZR1444400). The authors thank Dr. Suzanne Noble at UCSF for providing us the *C. albicans* homozygous gene deletion library and related plasmids as gift, and excellent supervision in postdoctoral education and training. We gratefully acknowledge Dr. Aaron P. Mitchell at the University of Georgia for valuable comments and helpful discussions. And we thank all the lab members in the Chen-lab at Institut Pasteur of Shanghai, Chinese Academy of Sciences and the members in the Liang-lab at State Key Laboratory of Trauma, Burns and Combined Injury, Third Military Medical University, for their help in discussion and preparation of the manuscript.

## Author contribution

CC and HL conceived and designed the study; CC, HL, YW and YM performed data analysis and wrote the manuscript; YW, YM, XH, TJ, YZ, CX, ZZ, LT, XM and XW conducted all the experiments; XC performed the statistical analysis of the data; KY performed the Drosophila infection; CC, HL, YW, YM and LP discussed the experiments and results.

## Supplementary Materials

**Table S1.** Strains used in this study.

**Table S2.** Plasmids used in this study.

**Table S3.** Primers used in this study.

### Figure supplement(s).

**Figure 1—figure supplement 1.**
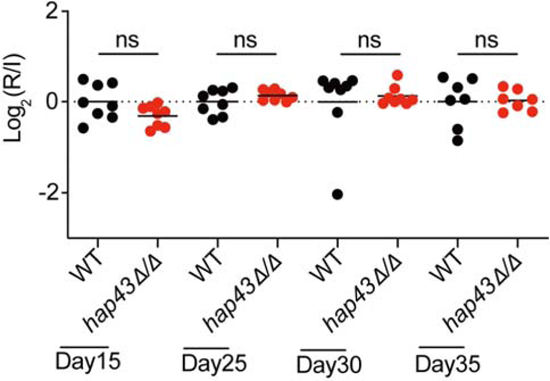
Mutant lacking *HAP43* exhibits no change in commensal fitness in NFD-treated mice. Mice (n=8) fed a normal Fe diet (NFD) were inoculated by gavage with 1:1 mixtures of the wild-type (WT) and *hap43Δ/Δ* mutant cells (1×10^8^ CFU per mice). The fitness value for each strain was calculated as the log_2_ ratio of its relative abundance in the recovered pool from the host (R) to the initial inoculum (I), and was determined by qPCR using strain-specific primers that could distinguish one from another. ns, no significance; by unpaired Student’s *t*-test.

**Figure 3—figure supplement 1.**
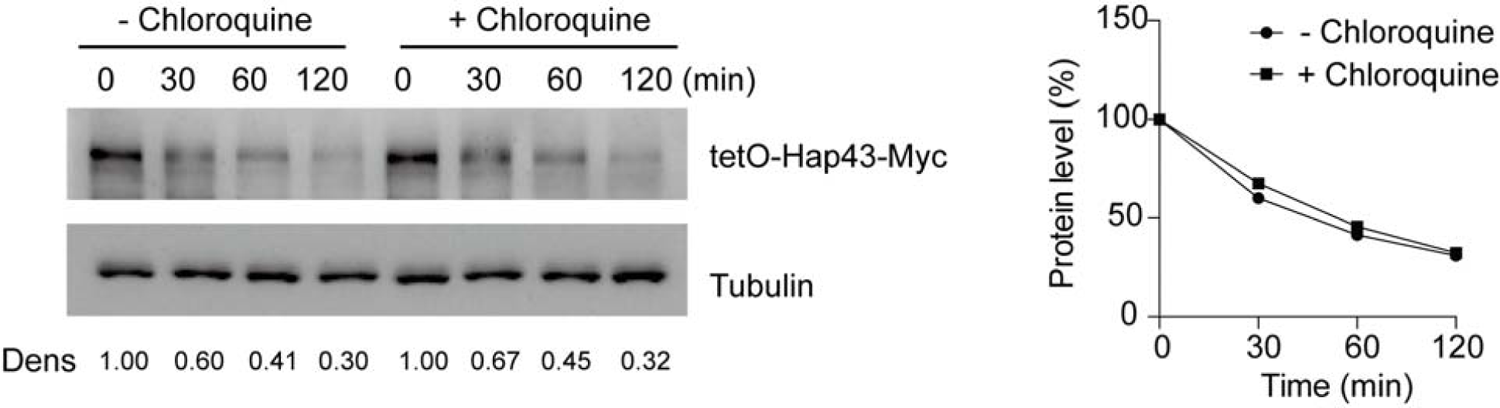
Chloroquine had no effect on Hap43 degradation under high iron conditions. After treatment with doxycycline, the WT cells stably expressing doxycycline-inducible Myc-tagged Hap43 (TetO-Hap43-Myc) were harvested, washed and treated with or without the lysosomal protease inhibitor chloroquine (100 mM). The turnover of Hap43-Myc in WT cells was evaluated through time-course experiments. Right panel: Hap43-Myc quantification after intensity analysis using Image J. For raw blots see **Figure 3—figure supplement 1—source data**.

**Figure 3—figure supplement 2.**
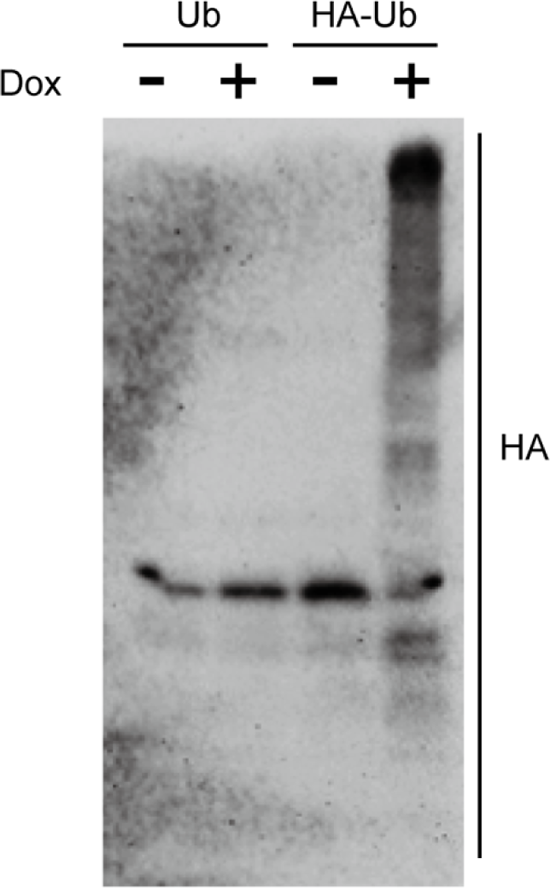
Immunoblots showing the induction of an epitope-tagged 3xHA-ubiquitin under the control of the doxycycline (DOX) inducible promoter. *C. albicans* cells co-expressing 3xHA-tagged ubiquitin and Hap43-Myc as well as Hap43-Myc cells were incubated in YPD supplemented with 50 μg/ml doxycycline (Dox) for 6 h. Log-phase cells were collected and lysed, followed by immunoblots of whole cell extracts with anti-HA antibodies. For raw blots see **Figure 3—figure supplement 2—source data**.

**Figure 4—figure supplement 1.**
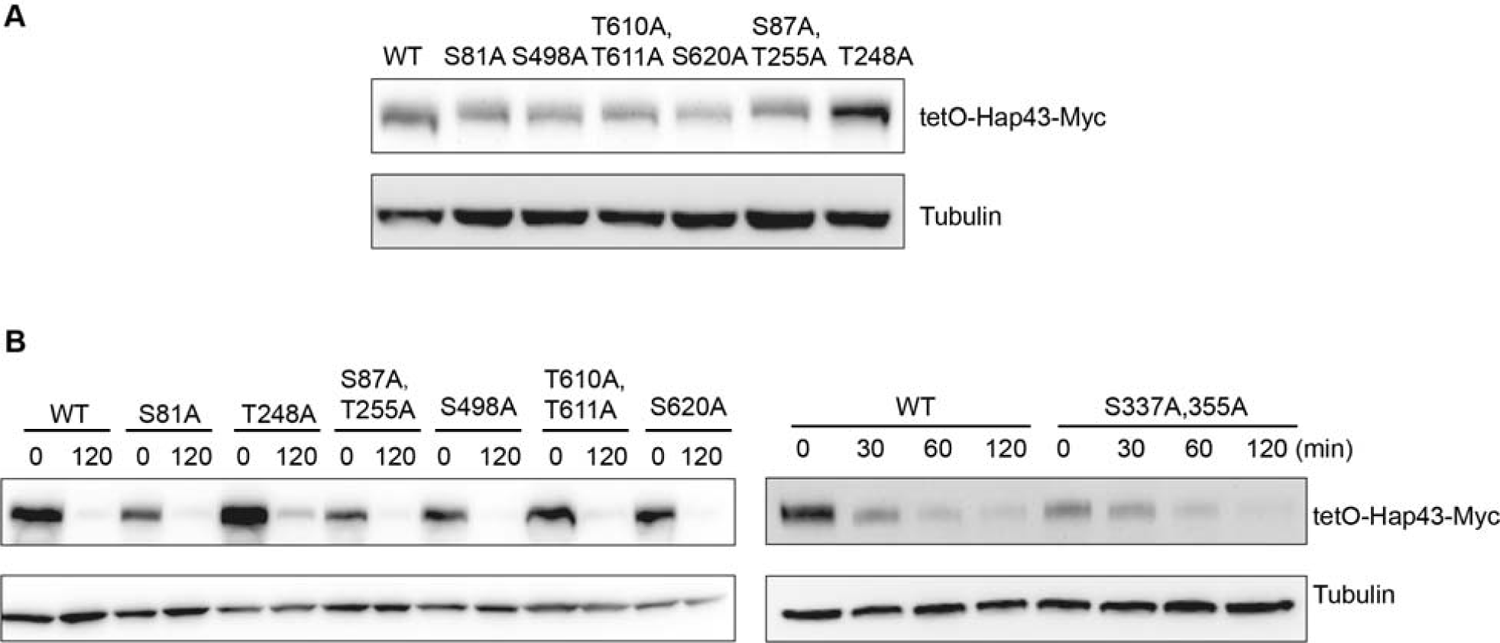
The Hap43 mutants harboring serine/threonine-to-alanine substitutions in its one or two putative phosphorylation sites showed the WT-like degradation patterns of Hap43 under high iron conditions. (**A)** Immunoblots of Hap43-Myc in strains expressing the indicated amino acid substitution allele of Hap43. Cells were treated at high iron conditions. **(B)** Strains expressing the indicated amino acid substitution allele of Hap43 under control of the inducible tetO promoter were treated with doxycycline. Cells were harvested in the exponential phase of growth, washed to remove doxycycline, and resuspended in fresh iron-replete medium. The turnover of Hap43-Myc in WT or relative mutant cells was then evaluated at 2 h following the tetO promoter shut-off by removal of doxycycline. For raw blots see **Figure 4—figure supplement 1—source data**.

**Figure 4—figure supplement 2.**
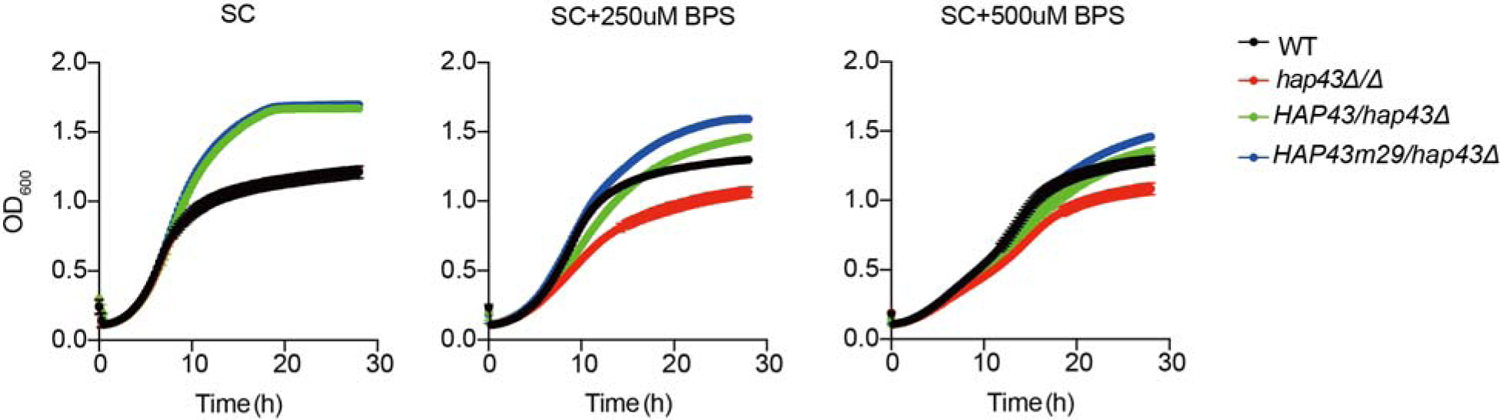
The mutants harboring amino acid substitutions or fragment truncation showed no defects in vegetative growth. Growth curve analysis of *HAP43* mutant strain harboring 29-point mutations in YPD liquid medium supplemented with 250 μM or 500 μM the impermeable iron chelator bathophenanthroline disulfonate (BPS) at 30 °C. OD_600_ readings were obtained every 15 min in a BioTek^TM^ Synergy^TM^ 2 Multi-mode Microplate Reader.

**Figure 4—figure supplement 3.**
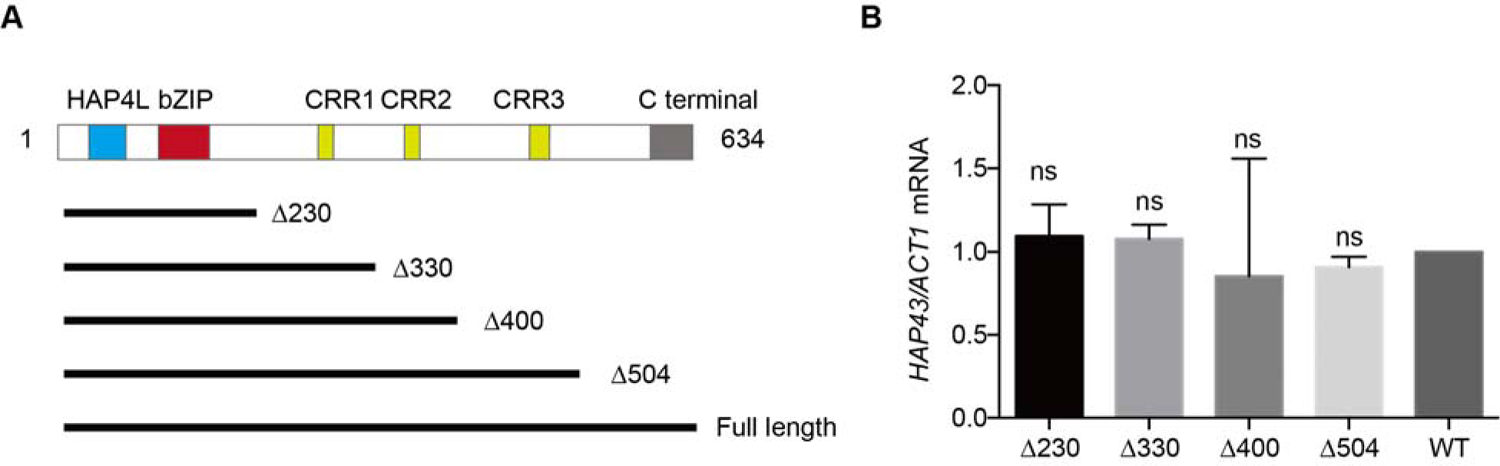
The critical phosphorylation region is essential for Hap43 stabilization. **(A)** Schematic diagram illustrating the Hap43 truncation proteins used as part of this study. The positions of the major domains identified in individuals with Hap43 are indicated. Numbers indicate the positions of the first and last amino acids, relative to the full-length protein. **(B)** qRT-PCR analysis for *HAP43* mRNA in WT and truncation mutant strains grown under iron-replete conditions. Transcript levels were normalized to the level of *ACT1* mRNA. Results from three independent experiments are shown. All data shown are means ± SD. ns, no significance; compared to WT, by one-way ANOVA with Dunnett’s test (B).

**Figure 4—figure supplement 4.**
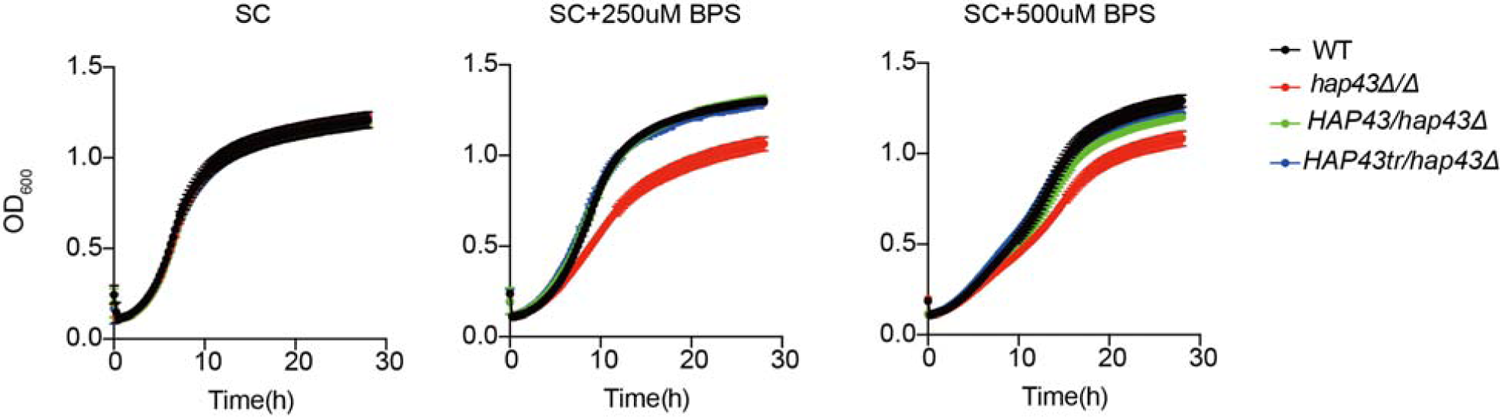
The mutants harboring amino acid substitutions or fragment truncation showed no defects in vegetative growth. Growth curve analysis of Hap43 truncation in YPD liquid medium supplemented with 250 μM or 500 μM the impermeable iron chelator bathophenanthroline disulfonate (BPS) at 30 °C. OD_600_ readings were obtained every 15 min in a BioTek^TM^ Synergy^TM^ 2 Multi-mode Microplate Reader.

**Figure 4—figure supplement 5.**
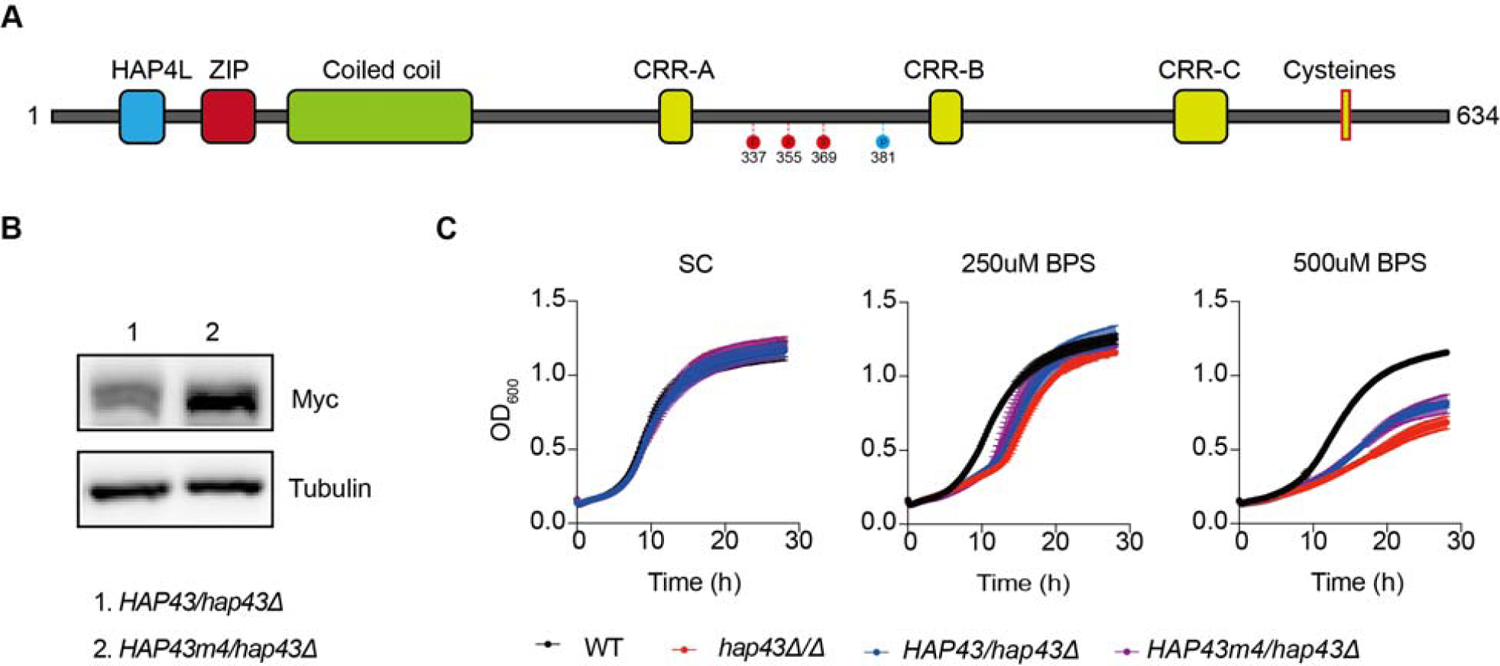
Identification of potential phosphorylation sites of Hap43. **(A)** Schematic representation of generating the *C. albicans* Hap43 mutation-4 (*HAP43*m4). Four putative S/T phosphorylation sites (S337/S355/S369/T381) in Hap43 were individually replaced with alanine residues. **(B)** Immunoblots of Hap43-Myc in strains expressing either the WT or the amino acid mutation (*HAP43*m4). Cells were treated at high iron conditions. **(C)** The *HAP43* mutant-4 strain (*HAP43m4/hap43Δ*) harboring amino acid substitutions showed no defects in vegetative growth. Growth curve analysis of indicated strains in SC liquid medium supplemented with 250 uM or 500 uM the impermeable iron chelator bathophenanthroline disulfonate (BPS) at 30 °C. OD_600_ readings were obtained every 15 min in a BioTek^TM^ Synergy^TM^ 2 Multi-mode Microplate Reader. For raw blots see **Figure 4—figure supplement 5—source data**.

**Figure 5—figure supplement 1.**
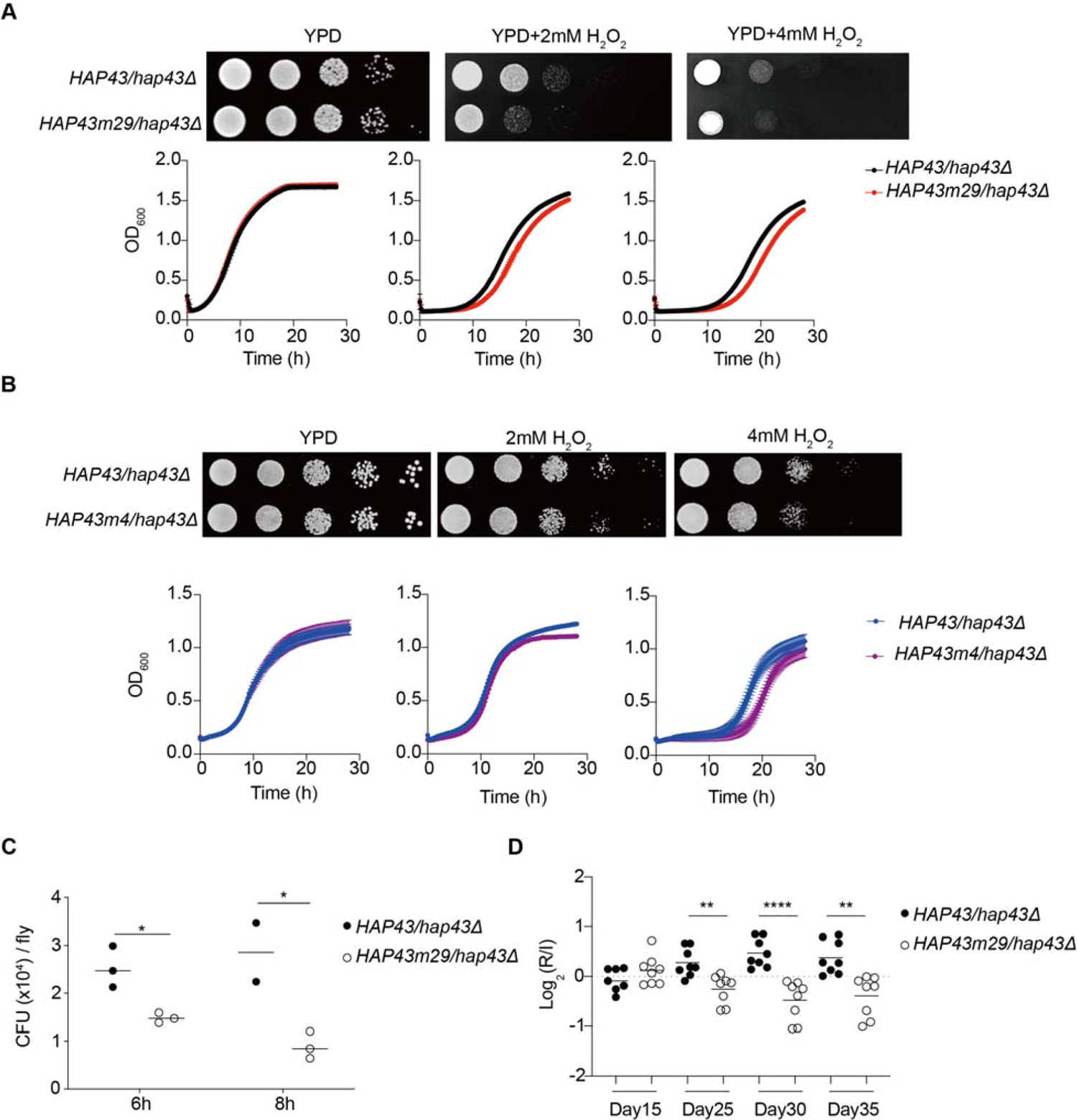
Hap43 phosphorylation is important for alleviating Fenton reaction-induced ROS toxicity and for GI colonization. Growth of *HAP43m29* mutant strain **(A)** or *HAP43m4* mutant strain **(B)** under oxidative stresses. The experiments were conducted the same way as describe in Figure. 5D. **(C)** Flies were fed on live yeast cells of indicated strains and the fungal burden (expressed as CFUs) were determined at different time points. **(D)** The *HAP43* mutant strain harboring 29-point mutations (*Hap43*m29) exhibits decreased commensal fitness in mice. Mice (n=8) were inoculated by gavage with 1:1 mixtures of WT and *Hap43m* mutant cells (1×10^8^ CFU per mice). The fitness value for each strain was calculated as the log_2_ ratio of its relative abundance in the recovered pool from the host (R) to the initial inoculum (I), and results from three independent experiments are shown. All data shown are means ± SD. \**p*< 0.05; \*\**p*<0.01; \*\*\*\**p*<0.0001; by two-way ANOVA with Sidak’s test **(C)** or unpaired Student’s *t*-test **(D)**.

**Figure 6—figure supplement 1.**
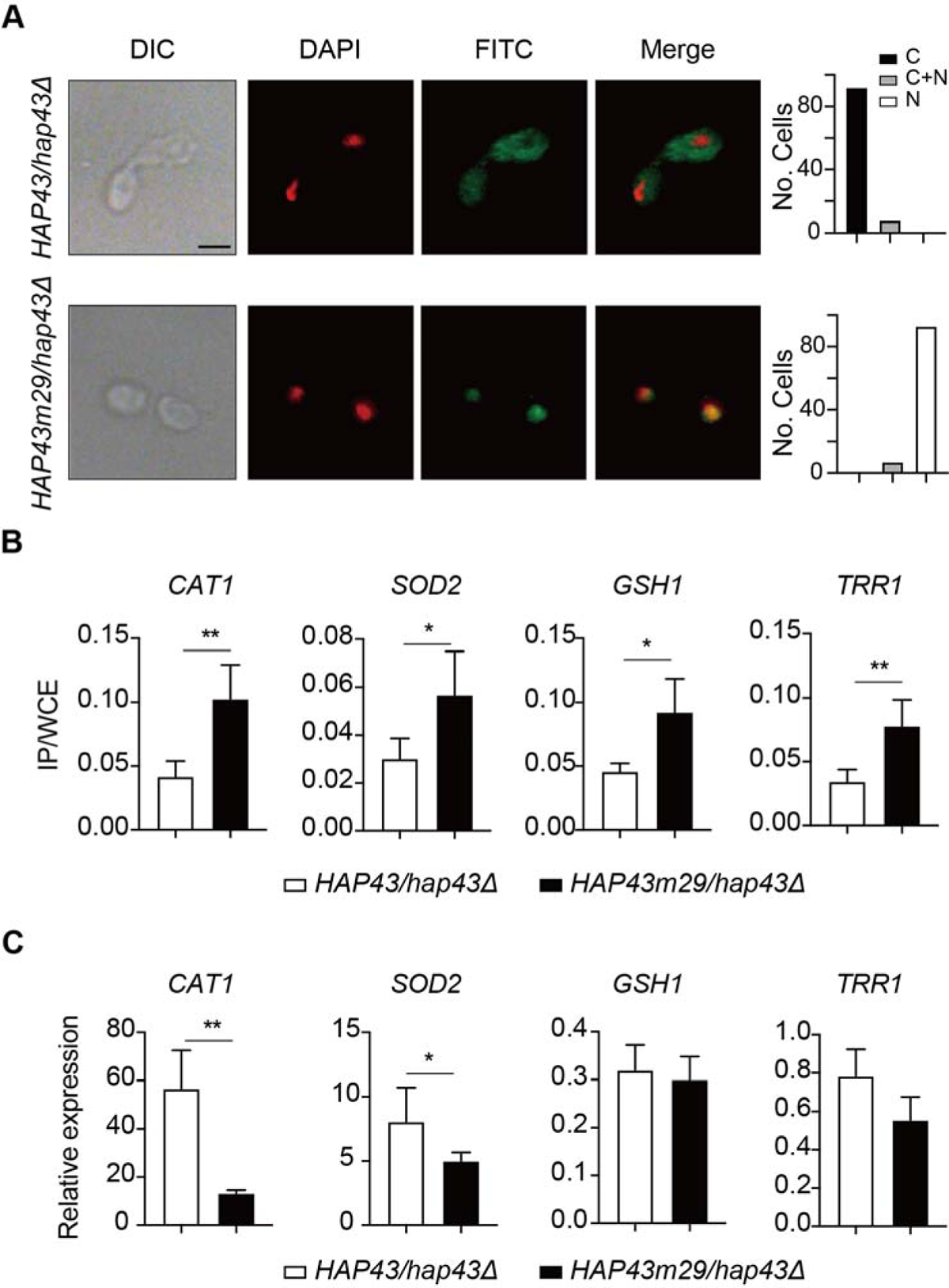
Iron-induced phosphorylation and degradation of Hap43 leads to de-repression of antioxidant genes. **(A)** Left panels: Indirect immunofluorescence of Hap43-Myc in *HAP43/hap43Δ* and *HAP43m29/hap43Δ* strains grown under iron-replete conditions. DIC represents phase images, DAPI represents nuclear staining, FITC represents Hap43-Myc staining, and Merge represents the overlay of Hap43-Myc and nuclear staining. Right panels: Quantification of the cellular distribution of Hap43. Each bar represents the analysis of at least 100 cells. C representing >90% cytoplasmic staining, N >90% nuclear staining, and C+N a mixture of cytoplasmic and nuclear staining. Scale bar, 5 µm. **(B)** ChIP of Hap43-Myc on the promoters that contain CCAAT boxes in a set of anti-oxidant genes. Overnight cultures of WT (*HAP43/hap43Δ*) and mutant harboring 29 substitutions (*HAP43m29/hap43Δ*) were grown and treated exactly the same way as described in Fig. 6b. **c** qRT-PCR analysis for mRNA levels of a set of antioxidant genes in WT (*HAP43/hap43Δ*) and mutant harboring 29 substitutions (*HAP43m29/hap43Δ*) grown under iron-replete conditions. Transcript levels were normalized to the level of *ACT1* mRNA. Results from three independent experiments are shown. All data shown are means ± SD. ns, no significance; \**p*< 0.05; \*\**p*<0.01; by unpaired Student’s *t*-test (B, C).

## References

Aguirre, J., Rios-Momberg, M., Hewitt, D., & Hansberg, W. (2005). Reactive oxygen species and development in microbial eukaryotes. Trends Microbiol, 13(3), 111–118. https://doi.org/10.1016/j.tim.2005.01.007

Andrews, N. C. (2008). Forging a field: the golden age of iron biology. Blood, 112(2), 219–230. https://doi.org/10.1182/blood-2007-12-077388

Avci, P., Freire, F., Banvolgyi, A., Mylonakis, E., Wikonkal, N. M., & Hamblin, M. R. (2016). Sodium ascorbate kills Candida albicans in vitro via iron-catalyzed Fenton reaction: importance of oxygenation and metabolism. Future Microbiol, 11, 1535–1547. https://doi.org/10.2217/fmb-2016-0063

Barber, M. F., & Elde, N. C. (2015). Buried Treasure: Evolutionary Perspectives on Microbial Iron Piracy. Trends Genet, 31(11), 627–636. https://doi.org/10.1016/j.tig.2015.09.001

Becker, D. M., Fikes, J. D., & Guarente, L. (1991). A cDNA encoding a human CCAAT-binding protein cloned by functional complementation in yeast. Proc Natl Acad Sci U S A, 88(5), 1968–1972. https://doi.org/10.1073/pnas.88.5.1968

Bink, A., Vandenbosch, D., Coenye, T., Nelis, H., Cammue, B. P., & Thevissen, K. (2011). Superoxide dismutases are involved in Candida albicans biofilm persistence against miconazole. Antimicrob Agents Chemother, 55(9), 4033–4037. https://doi.org/10.1128/AAC.00280-11

Bjorklund, S., & Gustafsson, C. M. (2005). The yeast Mediator complex and its regulation. Trends Biochem Sci, 30(5), 240–244. https://doi.org/10.1016/j.tibs.2005.03.008

Chen, C., & Noble, S. M. (2012). Post-transcriptional regulation of the Sef1 transcription factor controls the virulence of Candida albicans in its mammalian host. PLoS Pathog, 8(11), e1002956. https://doi.org/10.1371/journal.ppat.1002956

Chen, C., Pande, K., French, S. D., Tuch, B. B., & Noble, S. M. (2011). An iron homeostasis regulatory circuit with reciprocal roles in Candida albicans commensalism and pathogenesis. Cell Host Microbe, 10(2), 118–135. https://doi.org/10.1016/j.chom.2011.07.005

Chi, Y., Huddleston, M. J., Zhang, X., Young, R. A., Annan, R. S., Carr, S. A., & Deshaies, R. J. (2001). Negative regulation of Gcn4 and Msn2 transcription factors by Srb10 cyclin-dependent kinase. Genes Dev, 15(9), 1078–1092. https://doi.org/10.1101/gad.867501

Cohen, P. (2001). The role of protein phosphorylation in human health and disease. The Sir Hans Krebs Medal Lecture. Eur J Biochem, 268(19), 5001–5010. https://doi.org/10.1046/j.0014-2956.2001.02473.x

da Silva Dantas, A., Lee, K. K., Raziunaite, I., Schaefer, K., Wagener, J., Yadav, B., & Gow, N. A. (2016). Cell biology of Candida albicans-host interactions. Curr Opin Microbiol, 34, 111–118. https://doi.org/10.1016/j.mib.2016.08.006

Dantas Ada, S., Day, A., Ikeh, M., Kos, I., Achan, B., & Quinn, J. (2015). Oxidative stress responses in the human fungal pathogen, Candida albicans. Biomolecules, 5(1), 142–165. https://doi.org/10.3390/biom5010142

Dixon, S. J., & Stockwell, B. R. (2014). The role of iron and reactive oxygen species in cell death. Nat Chem Biol, 10(1), 9–17. https://doi.org/10.1038/nchembio.1416

Donko, A., Morand, S., Korzeniowska, A., Boudreau, H. E., Zana, M., Hunyady, L., Geiszt, M., & Leto, T. L. (2014). Hypothyroidism-associated missense mutation impairs NADPH oxidase activity and intracellular trafficking of Duox2. Free Radic Biol Med, 73, 190–200. https://doi.org/10.1016/j.freeradbiomed.2014.05.006

Dorn, A., Bollekens, J., Staub, A., Benoist, C., & Mathis, D. (1987). A multiplicity of CCAAT box-binding proteins. Cell, 50(6), 863–872. https://doi.org/10.1016/0092-8674(87)90513-7

El-Benna, J., Hurtado-Nedelec, M., Marzaioli, V., Marie, J. C., Gougerot-Pocidalo, M. A., & Dang, P. M. (2016). Priming of the neutrophil respiratory burst: role in host defense and inflammation. Immunol Rev, 273(1), 180–193. https://doi.org/10.1111/imr.12447

Frawley, E. R., Crouch, M. L., Bingham-Ramos, L. K., Robbins, H. F., Wang, W., Wright, G. D., & Fang, F. C. (2013). Iron and citrate export by a major facilitator superfamily pump regulates metabolism and stress resistance in Salmonella Typhimurium. Proc Natl Acad Sci U S A, 110(29), 12054–12059. https://doi.org/10.1073/pnas.1218274110

Galaris, D., Barbouti, A., & Pantopoulos, K. (2019). Iron homeostasis and oxidative stress: An intimate relationship. Biochim Biophys Acta Mol Cell Res, 1866(12), 118535. https://doi.org/10.1016/j.bbamcr.2019.118535

Ghosh, S., & Dass, J. F. P. (2016). Study of pathway cross-talk interactions with NF-kappaB leading to its activation via ubiquitination or phosphorylation: A brief review. Gene, 584(1), 97–109. https://doi.org/10.1016/j.gene.2016.03.008

Glittenberg, M. T., Kounatidis, I., Christensen, D., Kostov, M., Kimber, S., Roberts, I., & Ligoxygakis, P. (2011). Pathogen and host factors are needed to provoke a systemic host response to gastrointestinal infection of Drosophila larvae by Candida albicans. Dis Model Mech, 4(4), 515–525. https://doi.org/10.1242/dmm.006627

Grass, G., Otto, M., Fricke, B., Haney, C. J., Rensing, C., Nies, D. H., & Munkelt, D. (2005). FieF (YiiP) from Escherichia coli mediates decreased cellular accumulation of iron and relieves iron stress. Arch Microbiol, 183(1), 9–18. https://doi.org/10.1007/s00203-004-0739-4

Gupta, M., & Outten, C. E. (2020). Iron-sulfur cluster signaling: The common thread in fungal iron regulation. Curr Opin Chem Biol, 55, 189–201. https://doi.org/10.1016/j.cbpa.2020.02.008

He, X., Yu, J., Wang, M., Cheng, Y., Han, Y., Yang, S., Shi, G., Sun, L., Fang, Y., Gong, S. T., Wang, Z., Fu, Y. X., Pan, L., & Tang, H. (2017). Bap180/Baf180 is required to maintain homeostasis of intestinal innate immune response in Drosophila and mice. Nat Microbiol, 2, 17056. https://doi.org/10.1038/nmicrobiol.2017.56

Hershko, A., & Ciechanover, A. (1998). The ubiquitin system. Annu Rev Biochem, 67, 425–479. https://doi.org/10.1146/annurev.biochem.67.1.425

Hsu, P. C., Yang, C. Y., & Lan, C. Y. (2011). Candida albicans Hap43 is a repressor induced under low-iron conditions and is essential for iron-responsive transcriptional regulation and virulence. Eukaryot Cell, 10(2), 207–225. https://doi.org/10.1128/EC.00158-10

Jiang, X., Hoehenwarter, W., Scheel, D., & Lee, J. (2020). Phosphorylation of the CAMTA3 Transcription Factor Triggers Its Destabilization and Nuclear Export. Plant Physiol, 184(2), 1056–1071. https://doi.org/10.1104/pp.20.00795

Jung, W. H., Saikia, S., Hu, G., Wang, J., Fung, C. K., D’Souza, C., White, R., & Kronstad, J. W. (2010). HapX positively and negatively regulates the transcriptional response to iron deprivation in Cryptococcus neoformans. PLoS Pathog, 6(11), e1001209. https://doi.org/10.1371/journal.ppat.1001209

Kato, M. (2005). An overview of the CCAAT-box binding factor in filamentous fungi: assembly, nuclear translocation, and transcriptional enhancement. Biosci Biotechnol Biochem, 69(4), 663–672. https://doi.org/10.1271/bbb.69.663

Kobayashi, D., Kondo, K., Uehara, N., Otokozawa, S., Tsuji, N., Yagihashi, A., & Watanabe, N. (2002). Endogenous reactive oxygen species is an important mediator of miconazole antifungal effect. Antimicrob Agents Chemother, 46(10), 3113–3117. https://doi.org/10.1128/AAC.46.10.3113-3117.2002

Kumamoto, C. A., Gresnigt, M. S., & Hube, B. (2020). The gut, the bad and the harmless: Candida albicans as a commensal and opportunistic pathogen in the intestine. Curr Opin Microbiol, 56, 7–15. https://doi.org/10.1016/j.mib.2020.05.006

Lecker, S. H., Goldberg, A. L., & Mitch, W. E. (2006). Protein degradation by the ubiquitin-proteasome pathway in normal and disease states. J Am Soc Nephrol, 17(7), 1807–1819. https://doi.org/10.1681/ASN.2006010083

Lipford, J. R., & Deshaies, R. J. (2003). Diverse roles for ubiquitin-dependent proteolysis in transcriptional activation. Nat Cell Biol, 5(10), 845–850. https://doi.org/10.1038/ncb1003-845

Lu, Y., Su, C., Ray, S., Yuan, Y., & Liu, H. (2019). CO2 Signaling through the Ptc2-Ssn3 Axis Governs Sustained Hyphal Development of Candida albicans by Reducing Ume6 Phosphorylation and Degradation. mBio, 10(1). https://doi.org/10.1128/mBio.02320-18

Lund, E. K., Wharf, S. G., Fairweather-Tait, S. J., & Johnson, I. T. (1998). Increases in the concentrations of available iron in response to dietary iron supplementation are associated with changes in crypt cell proliferation in rat large intestine. J Nutr, 128(2), 175–179. https://doi.org/10.1093/jn/128.2.175

Lund, E. K., Wharf, S. G., Fairweather-Tait, S. J., & Johnson, I. T. (1999). Oral ferrous sulfate supplements increase the free radical-generating capacity of feces from healthy volunteers. Am J Clin Nutr, 69(2), 250–255. https://doi.org/10.1093/ajcn/69.2.250

Mahalhal, A., Williams, J. M., Johnson, S., Ellaby, N., Duckworth, C. A., Burkitt, M. D., Liu, X., Hold, G. L., Campbell, B. J., Pritchard, D. M., & Probert, C. S. (2018). Oral iron exacerbates colitis and influences the intestinal microbiome. PLoS One, 13(10), e0202460. https://doi.org/10.1371/journal.pone.0202460

Mao, Y., & Chen, C. (2019). The Hap Complex in Yeasts: Structure, Assembly Mode, and Gene Regulation. Front Microbiol, 10, 1645. https://doi.org/10.3389/fmicb.2019.01645

McNabb, D. S., & Pinto, I. (2005). Assembly of the Hap2p/Hap3p/Hap4p/Hap5p-DNA complex in Saccharomyces cerevisiae. Eukaryot Cell, 4(11), 1829–1839. https://doi.org/10.1128/EC.4.11.1829-1839.2005

Mello, E. O., Ribeiro, S. F., Carvalho, A. O., Santos, I. S., Da Cunha, M., Santa-Catarina, C., & Gomes, V. M. (2011). Antifungal activity of PvD1 defensin involves plasma membrane permeabilization, inhibition of medium acidification, and induction of ROS in fungi cells. Curr Microbiol, 62(4), 1209–1217. https://doi.org/10.1007/s00284-010-9847-3

Miret, S., Simpson, R. J., & McKie, A. T. (2003). Physiology and molecular biology of dietary iron absorption. Annu Rev Nutr, 23, 283–301. https://doi.org/10.1146/annurev.nutr.23.011702.073139

Muratani, M., & Tansey, W. P. (2003). How the ubiquitin-proteasome system controls transcription. Nat Rev Mol Cell Biol, 4(3), 192–201. https://doi.org/10.1038/nrm1049

Nelson, C., Goto, S., Lund, K., Hung, W., & Sadowski, I. (2003). Srb10/Cdk8 regulates yeast filamentous growth by phosphorylating the transcription factor Ste12. Nature, 421(6919), 187–190. https://doi.org/10.1038/nature01243

Nobile, C. J., Nett, J. E., Hernday, A. D., Homann, O. R., Deneault, J. S., Nantel, A., Andes, D. R., Johnson, A. D., & Mitchell, A. P. (2009). Biofilm matrix regulation by Candida albicans Zap1. PLoS Biol, 7(6), e1000133. https://doi.org/10.1371/journal.pbio.1000133

Noble, S. M., & Johnson, A. D. (2007). Genetics of Candida albicans, a diploid human fungal pathogen. Annu Rev Genet, 41, 193–211. https://doi.org/10.1146/annurev.genet.41.042007.170146

Olsen, J. V., Blagoev, B., Gnad, F., Macek, B., Kumar, C., Mortensen, P., & Mann, M. (2006). Global, in vivo, and site-specific phosphorylation dynamics in signaling networks. Cell, 127(3), 635–648. https://doi.org/10.1016/j.cell.2006.09.026

Pierre, J. L., Fontecave, M., & Crichton, R. R. (2002). Chemistry for an essential biological process: the reduction of ferric iron. Biometals, 15(4), 341–346. https://doi.org/10.1023/a:1020259021641

Pinkham, J. L., & Guarente, L. (1985). Cloning and molecular analysis of the HAP2 locus: a global regulator of respiratory genes in Saccharomyces cerevisiae. Mol Cell Biol, 5(12), 3410–3416. https://doi.org/10.1128/mcb.5.12.3410-3416.1985

Pizarro, F., Amar, M., & Stekel, A. (1987). Determination of iron in stools as a method to monitor consumption of iron-fortified products in infants. Am J Clin Nutr, 45(2), 484–487. https://doi.org/10.1093/ajcn/45.2.484

Punga, T., Bengoechea-Alonso, M. T., & Ericsson, J. (2006). Phosphorylation and ubiquitination of the transcription factor sterol regulatory element-binding protein-1 in response to DNA binding. J Biol Chem, 281(35), 25278–25286. https://doi.org/10.1074/jbc.M604983200

Puri, S., Kumar, R., Rojas, I. G., Salvatori, O., & Edgerton, M. (2019). Iron Chelator Deferasirox Reduces Candida albicans Invasion of Oral Epithelial Cells and Infection Levels in Murine Oropharyngeal Candidiasis. Antimicrob Agents Chemother, 63(4). https://doi.org/10.1128/AAC.02152-18

Reed, C., Ibrahim, A., Edwards, J. E., Jr., Walot, I., & Spellberg, B. (2006). Deferasirox, an iron-chelating agent, as salvage therapy for rhinocerebral mucormycosis. Antimicrob Agents Chemother, 50(11), 3968–3969. https://doi.org/10.1128/AAC.01065-06

Ren, T., Zhu, H., Tian, L., Yu, Q., & Li, M. (2020). Candida albicans infection disturbs the redox homeostasis system and induces reactive oxygen species accumulation for epithelial cell death. FEMS Yeast Res, 20(4). https://doi.org/10.1093/femsyr/foz081

Ryan, T. P., & Aust, S. D. (1992). The role of iron in oxygen-mediated toxicities. Crit Rev Toxicol, 22(2), 119–141. https://doi.org/10.3109/10408449209146308

Schieber, M., & Chandel, N. S. (2014). ROS function in redox signaling and oxidative stress. Curr Biol, 24(10), R453–462. https://doi.org/10.1016/j.cub.2014.03.034

Schwartz, A. J., Das, N. K., Ramakrishnan, S. K., Jain, C., Jurkovic, M. T., Wu, J., Nemeth, E., Lakhal-Littleton, S., Colacino, J. A., & Shah, Y. M. (2019). Hepatic hepcidin/intestinal HIF-2alpha axis maintains iron absorption during iron deficiency and overload. J Clin Invest, 129(1), 336–348. https://doi.org/10.1172/JCI122359

Singh, A., Kaur, N., & Kosman, D. J. (2007). The metalloreductase Fre6p in Fe-efflux from the yeast vacuole. J Biol Chem, 282(39), 28619–28626. https://doi.org/10.1074/jbc.M703398200

Singh, R. P., Prasad, H. K., Sinha, I., Agarwal, N., & Natarajan, K. (2011). Cap2-HAP complex is a critical transcriptional regulator that has dual but contrasting roles in regulation of iron homeostasis in Candida albicans. J Biol Chem, 286(28), 25154–25170. https://doi.org/10.1074/jbc.M111.233569

Skrahina, V., Brock, M., Hube, B., & Brunke, S. (2017). Candida albicans Hap43 Domains Are Required under Iron Starvation but Not Excess. Front Microbiol, 8, 2388. https://doi.org/10.3389/fmicb.2017.02388

Swaney, D. L., Beltrao, P., Starita, L., Guo, A., Rush, J., Fields, S., Krogan, N. J., & Villen, J. (2013). Global analysis of phosphorylation and ubiquitylation cross-talk in protein degradation. Nat Methods, 10(7), 676–682. https://doi.org/10.1038/nmeth.2519

Symeonidis, A. S. (2009). The role of iron and iron chelators in zygomycosis. Clin Microbiol Infect, 15 *Suppl 5*, 26–32. https://doi.org/10.1111/j.1469-0691.2009.02976.x

Thon, M., Al Abdallah, Q., Hortschansky, P., Scharf, D. H., Eisendle, M., Haas, H., & Brakhage, A. A. (2010). The CCAAT-binding complex coordinates the oxidative stress response in eukaryotes. Nucleic Acids Res, 38(4), 1098–1113. https://doi.org/10.1093/nar/gkp1091

Thrower, J. S., Hoffman, L., Rechsteiner, M., & Pickart, C. M. (2000). Recognition of the polyubiquitin proteolytic signal. EMBO J, 19(1), 94–102. https://doi.org/10.1093/emboj/19.1.94

Tumusiime, S., Zhang, C., Overstreet, M. S., & Liu, Z. (2011). Differential regulation of transcription factors Stp1 and Stp2 in the Ssy1-Ptr3-Ssy5 amino acid sensing pathway. J Biol Chem, 286(6), 4620–4631. https://doi.org/10.1074/jbc.M110.195313

Valko, M., Morris, H., Mazur, M., Rapta, P., & Bilton, R. F. (2001). Oxygen free radical generating mechanisms in the colon: do the semiquinones of vitamin K play a role in the aetiology of colon cancer? Biochim Biophys Acta, 1527(3), 161–166. https://doi.org/10.1016/s0304-4165(01)00163-5

Van Breusegem, F., Vranova, E., Dat, J. F., & Inze, D. (2001). The role of active oxygen species in plant signal transduction. Plant Science, 161(3), 405–414. Doi https://doi.org/10.1016/S0168-9452(01)00452-6

von Hagen, J., & Michelsen, U. (2013). Cellular fractionation--yeast cells. Methods Enzymol, 533, 31–39. https://doi.org/10.1016/B978-0-12-420067-8.00004-0

Wong, Y. H., Lee, T. Y., Liang, H. K., Huang, C. M., Wang, T. Y., Yang, Y. H., Chu, C. H., Huang, H. D., Ko, M. T., & Hwang, J. K. (2007). KinasePhos 2.0: a web server for identifying protein kinase-specific phosphorylation sites based on sequences and coupling patterns. Nucleic Acids Res, 35(Web Server issue), W588–594. https://doi.org/10.1093/nar/gkm322

